# Effects of Drought, Followed by Waterlogging, on Biomass, Leaf Physiology and Chemistry of Wheat Partly Depend on Fertilisation

**DOI:** 10.64898/2026.09.30.755601

**Authors:** Rabea Schweiger, Caroline Müller

## Abstract

Drought and waterlogging are prominent climate change challenges that, alone or in sequence, pronouncedly constrain agricultural crop production. A better understanding of relationships between water availability and nutrition regarding plant functioning is needed to optimise fertilisation under these challenges. We investigated impacts of drought, waterlogging and drought followed by waterlogging under two fertilisation regimes (low, normal) on various traits of wheat (*Triticum aestivum*). Next to biomass and photosynthesis measurements, cross-platform metabolomics was applied for in-depth metabolic analyses of flag leaves. We found detrimental effects of drought and drought-waterlogging on shoot and root biomass, especially at normal fertilisation, while photosynthetic traits were not or only slightly affected. Foliar nitrogen contents were higher in plants exposed to drought or to drought-waterlogging. Numerous primary and specialised metabolites were modulated by fertilisation and/ or watering. In particular, amino acids, benzoxazinoids and flavonoids were affected by fertilisation. Due to watering especially several amino acids changed in concentrations, with the induction pattern partly depending on fertilisation. Our study deepens the understanding of the interplay between water and nutrient availabilities as well as nutritional demands, growth and metabolism of wheat. This is highly relevant to balance sustainable agricultural nutrient management under the challenges of ongoing climate change.

## 1 Introduction

Progressive climate change poses several challenges for plants, including important crop species. Droughts as well as floods caused by heavy rainfalls are predicted to increase (IPCC 2023), with the latter leading to (partial) waterlogging of plants. Drought and waterlogging may also occur one after another. While both challenges have been shown to affect various traits of plants if investigated separately (Farooq et al. 2009; Zhang et al. 2025), impacts of drought-waterlogging cycles and underlying molecular mechanisms have not yet been extensively studied (Renziehausen et al. 2024). Moreover, adequate fertilisation may be decisive for the performance of agricultural crops under drought-waterlogging sequences.

Drought has been shown to affect plant biomass, resource allocation, physiology and metabolism. Crop yields are generally lower under drought (Dietz et al. 2021). Plants exposed to drought often invest more resources in roots than in shoots, with partly different investments in stems, leaves and reproductive structures (Eziz et al. 2017; Nezhadahmadi et al. 2013). Photosynthesis may be restricted under drought because stomata are closed to reduce water loss and/ or due to non-stomatal limitations; stomatal closure may be accompanied by an elevated instantaneous water use efficiency (WUE), i.e., a higher CO_2_ assimilation rate (A) per transpiration rate (E) (Flexas and Medrano 2002; Kang and Kang 2019). Drought-exposed bread wheat plants (*Triticum aestivum*, Poaceae) had higher nitrogen (N) contents in vegetative and reproductive aboveground plant parts compared to well-watered plants, while carbon (C) contents were less affected (Pons and Müller 2022). Due to drought, also various primary and specialised plant metabolites can be modulated in concentrations. For example, several amino acids, such as proline and γ-aminobutyric acid (GABA), commonly increase (Fàbregas and Fernie 2019; Szabados and Savouré 2010). In wheat, benzoxazinoids accumulated in flag leaves under pulsed drought, while some flavonoids had higher, others lower concentrations under drought compared to well-watered plants (Stallmann et al. 2020).

Similar to drought, waterlogging of plants or plant parts has far-reaching effects on plant functions. However, some of these are mediated by hypoxic stress due to oxygen deficiency in the submerged plant parts. Waterlogging leads to the formation of aerenchyma or adventitious roots, the induction of anaerobic metabolism or tissue decay in roots (Geng et al. 2026; Voesenek and Bailey-Serres 2015; Zhang et al. 2025). Moreover, several systemic effects of waterlogging have been described, some of which may be related to dysfunctional roots impairing water uptake (Renziehausen et al. 2024). As under drought, yields and photosynthesis are reduced under waterlogging in various crop species (Tian et al. 2021). In wheat, the foliar concentrations of N and other elements were lower in waterlogged compared to control plants and gene expression and metabolism of both roots and leaves were affected by waterlogging of the roots (Cid et al. 2024).

Plant responses to drought followed by waterlogging have been less well investigated yet and may be complex, as these challenges can have opposing effects or enhance each other (Renziehausen et al. 2024). In wheat, photosynthetic CO_2_ assimilation was impaired in plant subjected to drought and subsequent waterlogging (Saha and Johnson 2025). Contrarily, in another study on wheat, neither photosynthesis nor the WUE of flag leaves were affected by drought or by drought followed by waterlogging, while plants showed less biomass and various metabolic changes in leaf transcripts and metabolites at the end of these phases compared to well-watered plants (Schweiger et al. 2023). In maize (*Zea mays*, Poaceae), negative effects of drought and of drought followed by waterlogging on biomass- and photosynthesis-related traits were accompanied by different metabolic changes (Keya et al. 2025).

Together with water, nutrient availability is considered to be highly limiting for agricultural production, both playing a crucial role for plant growth and development (Gonzalez-Dugo et al. 2010). Nutrients may also affect metabolism, as shown for wheat, where the leaf metabolic composition was shifted depending on N fertilisation (Heyneke et al. 2017; Zhang et al. 2017). Under drought and waterlogging, nutrient deficiencies induced by these challenges probably largely contribute to yield losses (Arif et al. 2025; Swain et al. 2023). Such nutrient deficiencies and other negative effects of drought and waterlogging may be partly compensated by additional fertilisation (Drobnitch et al. 2025; Xu et al. 2026; Zhang et al. 2025). However, fertiliser application imposes a trade-off between agricultural production and global food security as well as environmental sustainability due to fertiliser-induced water pollution and greenhouse gas emissions (Penuelas et al. 2023). Hence, (further) reductions in fertiliser application and nutrient losses are required until 2030 according to the Farm to Fork Strategy (European Commission 2020). In that context, more studies are needed that investigate how different fertilisation rates affect the responses of crops to drought and drought followed by waterlogging.

The aim of this study was to elucidate the role of fertilisation in the responses of the important crop species *T. aestivum* to drought, waterlogging and to a drought-waterlogging sequence. Various plant traits, including biomass-as well as photosynthesis-related variables and numerous metabolites, measured via multi-omics on different analytical platforms, were assessed. We hypothesised that well-watered plants under normal fertilisation show a higher shoot and root biomass, shoot-to-root ratio, harvest index, CO_2_ assimilation rate and foliar N content than plants under low fertilisation, along with various leaf metabolic differences between these groups. We expected that drought-treated plants have a lower shoot and root biomass, shoot-to-root ratio and CO_2_ assimilation rate, but a higher WUE and leaf N content than well-watered plants. We assumed an accumulation of amino acids and benzoxazinoids and shifts in the concentrations of flavonoids and other metabolites under drought. For plants subjected to waterlogging, we proposed a lower shoot and root biomass, a higher shoot-to-root ratio as well as a lower CO_2_ assimilation rate and foliar N content than in well-watered plants, accompanied by various metabolic effects. For plants that experienced a drought-waterlogging sequence, we expected more pronounced effects compared to plants exposed to single challenges, but no effects on the shoot-to-root-ratio and foliar N content due to presumably opposite impacts by drought and waterlogging on these traits. We assumed that the effects of challenging water regimes on biomass, photosynthesis and foliar N content are more pronounced under low compared to normal fertilisation, while the metabolic responsiveness (i.e., the number of modulated metabolites) is higher under normal fertilisation.

## 2 Materials and Methods

### 2.1 Plant Materials and Treatments

The design was similar to a previous experiment (Schweiger et al. 2023), but we now included a differential fertilisation treatment and performed additional chemical analyses. The experiment was conducted in a climate chamber (20°C, relative humidity 70%, 16:8 h light:dark, light from 7 AM to 11 PM). One hundred pots (2 L; 11.3 x 11.3 x 21.5 cm; with holes at bottom, placed on individual dishes) were filled with a wet 1:2 (v:v) mixture of soil (Fruhstorfer special substrate type P, Hawita Group, Vechta, Germany) and river sand, which had been steamed (≥ 90°C) before. The initial SWC (substrate water content, i.e., mass of water per substrate dry mass) was determined with the gravimetric method (ISO 11465 1993), using four substrate subsamples, and was 28.6%. Grains of bread wheat (*Triticum aestivum*, spring cultivar Tybalt, W. von Borries-Eckendorf, Leopoldshöhe, Germany) were sown in two opposite corners of the pots with 10 cm distance to each other, with two grains in each corner (1 cm distance); if more than one seedling per corner emerged, excess seedlings were removed from seven days post sowing (dps) onwards, yielding pots with two seedlings in opposite corners. For an initial establishment phase, all pots were watered to a SWC of 26.8% from nine dps onwards every second day (with some exceptions, see Figure 1), by weighing randomly chosen pots and watering all pots with tap water according to the mean weight loss.

**FIGURE 1.**
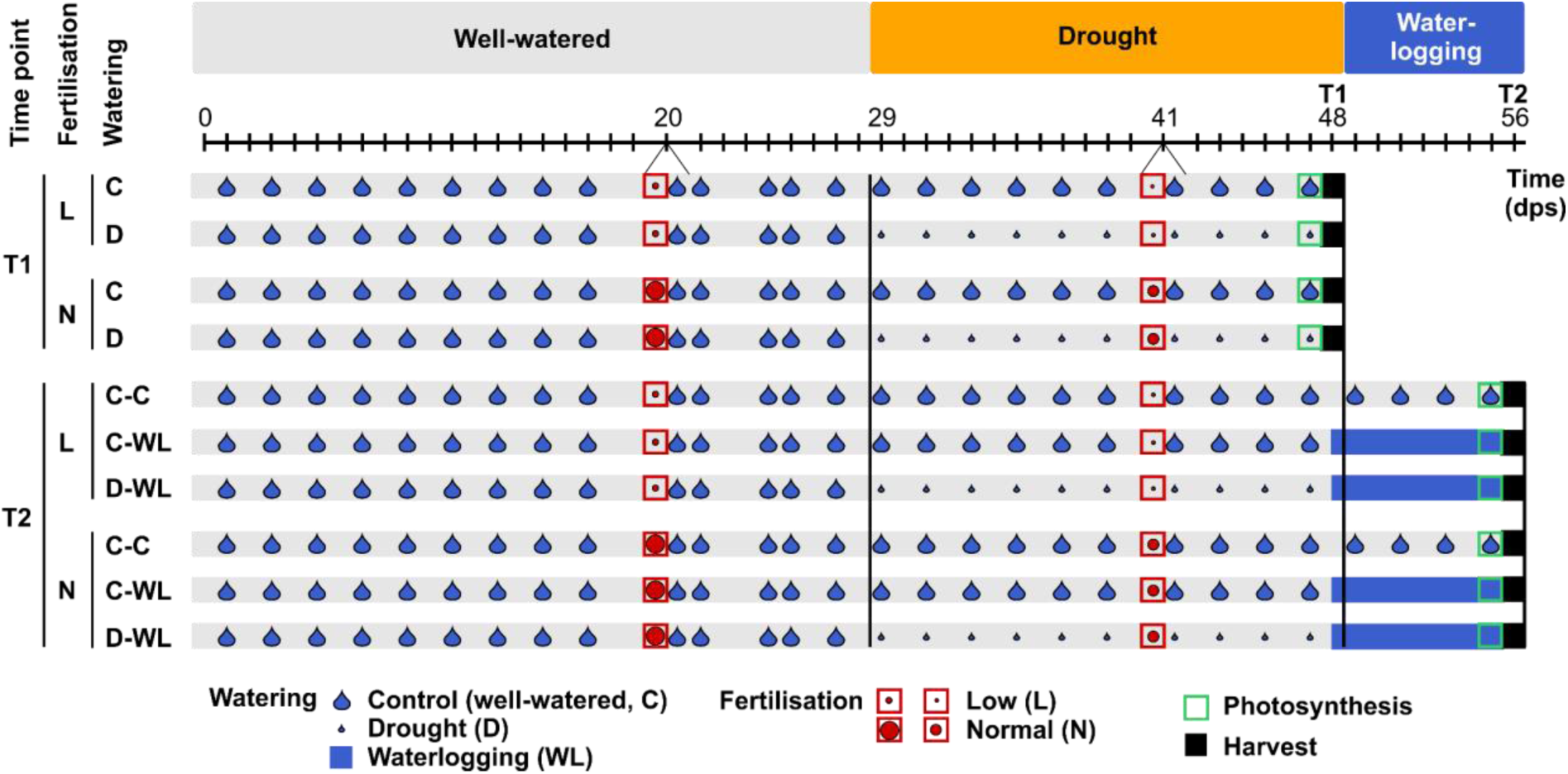
Timeline of treatments of *Triticum aestivum* plants of various groups. Plants were subjected to different water (C, well-watered; D, drought; WL, waterlogging) and fertilisation (L, low; N, normal) regimes. Four treatment groups were harvested at the end of the drought phase (T1), while the other six groups were harvested at the end of the subsequent waterlogging phase (T2). One day before harvest, photosynthesis was measured; *n*=10 biological replicates per group; dps, days post sowing.

At 29 dps, plants were randomly assigned to one of ten treatment (watering x fertilisation) groups, with *n*=10 pots each (Figure 1), using two subsequent different watering phases with two levels (drought phase: control, drought) or three levels (waterlogging phase: control all time, control followed by waterlogging, drought followed by waterlogging) and two fertilisation regimes (low and normal). A randomised block design was used, changing the positions of pots within blocks regularly. From 29 dps on, representative subsets of pots per treatment group were used to determine the watering volumes gravimetrically. In the drought phase (29-48 dps), control pots (C) were kept well-watered at 26.8% SWC, while pots of the drought treatment (D) were not watered until their mean SWC had fallen below 8.9% SWC and were then watered to 8.9% SWC every other day. After this phase (48 dps), a subset of plants was harvested (see 2.3; T1, four treatment groups with *n*=10 per group, Figure 1). In the subsequent waterlogging phase (48-56 dps; six treatment groups), control pots (C-C) were watered to 26.8% SWC and placed in empty canisters (5 L, Purania Stilles Quellwasser, TSI Consumer Goods, Zeven, Germany; cut at the top), while water-logged pots (WL), previously kept under control (C-WL) or drought conditions (D-WL), were placed in canisters filled with water up to few (< 5) cm above the substrate level and filled back up to that level every other day. Each of the water treatments was performed under low (L) and normal (N) fertilisation. Pots were fertilised with Plantosan N-P_2_O_5_-K_2_O 20-10-15 (containing 6% MgO, 2% S and traces of B, Cu, Fe, Mn, Mo, Zn; Manna, Düsseldorf, Germany) at 20 and 41 dps before watering the plants, with 0.67 g (20 dps) and 0.4 g (41 dps) per pot for the low fertilisation and 2 g (20 dps) and 1.2 g (41 pds) per pot for the normal fertilisation treatment. Plants were stabilised with bast cord at wooden sticks, fixed to the outer pot side at 19 dps and sticks removed before placing the plants into the canisters. At 56 dps, the second subset of plants was harvested (T2, six treatment groups with *n*=10 per group).

### 2.2 Measurements of Chlorophyll Fluorescence and Gas Exchange

Photosynthesis measurements were conducted one day before harvesting the two subsets of plants (T1: 47 dps; T2: 55 dps, only *n*=5-6 replicates per group due to time constraints) between 1-8 PM at one plant per pot close to the middle of the flag leaves of the main stems. A pulse-amplitude-modulated photosynthesis yield analyser (MINI-PAM, Walz, Effeltrich, Germany) was used to assess the effective photochemical quantum yield of photosystem II (ΔF/F_m_’) at one leaf position. At the adjacent leaf position, gas exchange [CO_2_ assimilation rate A, transpiration rate E; instantaneous water use efficiency WUE = A/E (Kang and Kang 2019)] was measured with a GFS-3000 (Walz), with a flow of 550 µmol s^-1^ and with 20°C, 415 ppm CO_2_ and 70% relative humidity in the leaf cuvette. For all measurements, light intensities close to the ones measured at leaf height (photosynthetic photon flux densities for MINI-PAM and GFS, respectively: 304 and 300 µmol m^-2^ s^-1^ at T1; 342 and 345 µmol m^-2^ s^-1^ at T2) were set. Plants were watered after these measurements.

### 2.3 Harvests, Carbon and Nitrogen Analyses

Harvests were performed between 1 PM and 5 PM. For chemical analyses, two leaf blades (flag leaf and next older leaf) from the main stem were separately harvested, using the plant per pot that had not been taken for photosynthesis measurements. Leaves were shock-frozen in liquid N_2_, stored at −80°C, freeze-dried and weighed. The total remaining aboveground plant material of both plants per pot was cut, separated in vegetative and generative parts (at T2 only, as at T1 ears had not emerged yet), dried (60°C) and weighed. The total shoot dry biomass was derived as the sum of that of the separately harvested leaf blades and remaining plant parts. Pots were stored at −20°C for few days and then roots washed from the substrate at room temperature, dried at 60°C and weighed. Dry biomass-based shoot-to-root ratios and (for T2, when emerged ears were found in all pots) harvest indices, i.e., the ratio of ear biomass to total shoot biomass, were determined.

The leaf blades were homogenised and leaf powder was split for different chemical analyses, using the flag leaf for analyses of carbon and nitrogen content and amino acids (see 2.4) and the next older leaf for other chemical analyses (see 2.5, 2.6). Carbon and nitrogen contents were measured with a Vario MICRO Cube (Elementar Analysensysteme, Hanau, Germany).

### 2.4 Analysis of Amino Acids

For amino acid analysis, leaf powder was extracted thrice with 80% (v:v) methanol (LC-MS grade, Fisher Scientific, Loughborough, UK) containing the internal standards L-norvaline (for primary amino acids) and sarcosine (for secondary amino acids) (Agilent Technologies, Waldbronn, Germany), followed by filtering (syringe filters 0.2 µm, Phenomenex, Torrance, CA, USA), similarly as previously described (Schweiger et al. 2023). Extracts were analysed on a high performance liquid chromatograph equipped with a fluorescence detector (HPLC-FLD, abbreviated as LC-FLD; Vanquish Core, Thermo Fisher Scientific, Dreieich, Germany). Pre-column derivatisation took place in the autosampler at 6°C, using borate buffer (Agilent Technologies), ortho-phthaldialdehyde (OPA, Agilent Technologies), 9-fluorenyl-methyl chloroformate (FMOC, Agilent Technologies) and 1 M acetic acid (eluent additive for LC-MS, J&K Scientific, Pforzheim, Germany) as neutralisation reagent. Derivatised amino acids were separated at 40°C on an Accucore C18 column (150 x 3 mm with pre-column, 2.6 µm particle size, Thermo Fisher Scientific), with eluents A [1.4 g Na_2_HPO_4_ (AppliChem, Darmstadt, Germany), 3.8 g Na_2_B_4_O_7_ x 10 H_2_O (Sigma-Aldrich, Steinheim, Germany) and 32 mg NaN_3_ (Carl Roth, Karlsruhe, Germany) in 1 L deionised water, pH 7.8] and B [4.5:4.5:1 mixture of acetonitrile (LC-MS grade, Fisher Scientific), methanol and deionised water] at a flow of 0.6 mL min^-1^ with a gradient from 2% to 10% B within 0.4 min, to 34.5% B within 11.7 min, to 100%B within 10 min, followed by column cleaning and equilibration. Primary amino acids were detected at an excitation wavelength of 337 nm and an emission wavelength of 442 nm, while 260 nm (excitation) and 325 nm (emission) were used for secondary amino acids. Ten blanks (without plant material) and standard mixes were measured in addition. Amino acid data were analysed with Chromeleon 7.2.10 ES MUi (Thermo Fisher Scientific), with identifications based on comparison of retention times (RTs) with standards. Identified amino acids were quantified using peak areas divided by those of the corresponding internal standard and, after mean peak areas in blanks were subtracted, data were divided by the sample dry mass. Minute peaks of ornithine, lysine and hydroxyproline were excluded due to poor chromatographic behaviour.

### 2.5 Analysis of Further Primary Metabolites

For profiling of further primary metabolites such as carbohydrates and organic acids, leaf powder was extracted in a 1:2.5:1 (v:v:v) mixture of chloroform (HPLC grade, AppliChem), methanol and deionised water, including the internal standard phenyl-β-D-glucopyranoside (Sigma-Aldrich, St. Louis, MO, USA), similar as in Schweiger et al. (2023). Water was added for phase separation, subsamples of the water/ methanol phases were dried under nitrogen and samples derivatised at 37°C in two subsequent steps: methoximation with 20 mg mL^-1^ *O*-methylhydroxylamine hydrochloride (Alfa Aesar, Kandel, Germany) in pyridine (HPLC grade, Honeywell, Charlotte, NC, USA) for 90 min, followed by silylation with *N*-methyl-*N*-trimethylsilyl-trifluoroacetamide (Macherey-Nagel, Düren, Germany) for 30 min. Samples were then analysed on a gas chromatograph coupled to a mass spectrometer (GC-MS; GC-2010 Plus coupled to QP2020, Shimadzu, Kyoto, Japan). After injection (225°C, 1:10 split), chromatographic separation took place at a helium flow of 1.14 mL min^-1^ on a VF-5ms column (30 m, with 8 m guard column, inner diameter 0.25 mm, film thickness 0.25 µm, Agilent J&W GC Columns, Santa Clara, CA, USA), with the oven temperature being 80°C for 3 min, followed by an increase (5°C min^-1^) to 310°C (held for 2 min) and an increase (15°C min^-1^) to 325°C (held for 3 min). The transfer line and ion source temperature were set to 250°C and 230°C, respectively. Electron impact ionisation was applied at 70 eV, with detection of *m*/*z* (mass-to-charge) ratios of 40 to 600. Ten blanks and, for calculation of arithmetic retention indices (AI) (van den Dool and Kratz 1962), a series of *n*-alkanes (C_7_-C_40_, Sigma-Aldrich) were measured in addition. GC-MS data were analysed using GCMS Postrun Analysis 4.45 (Shimadzu). Analytes were identified using a laboratory-internal library, comparing AIs and mass spectra as well as characteristic *m*/*z* published in the Golm metabolome database (Hummel et al. 2010; Schauer et al. 2005). Metabolites were quantified based on peak heights of total ion count chromatograms, divided by the peak height of phenyl-β-D-glucopyranoside, blanks (averaged) were subtracted and peak heights divided by the sample dry mass. If metabolites were represented by more than one analyte, these were summed up. Minor peaks of shikimic acid, quinic acid and galactose were excluded due to suboptimal chromatographic behaviour.

### 2.6 Analysis of Specialised Metabolites

For metabolic fingerprinting, leaf powder was extracted with ice-cold 90% (v:v) methanol, containing the internal standard mefenamic acid (Sigma-Aldrich), in an ultrasonic bath for 15 min, followed by filtering (0.2 µm filters, Phenomenex), as in Schweiger et al. (2021). This extraction procedure yields (semi-)polar metabolites, mainly specialised metabolites. Extracts were subjected to an ultra-HPLC quadrupole time-of-flight mass spectrometer [UHPLC (Dionex UltiMate 3000, Thermo Fisher Scientific, San José, CA, USA) coupled to QTOF-MS/MS (compact, Bruker Daltonics, Bremen, Germany); abbreviated as LC-MS in the following]. The autosampler was set to 5°C. Metabolites were separated on a Kinetex XB-C18 column (150 x 2.1 mm with pre-column, 1.7 µm particle size, Phenomenex) at 45°C and a flow of 0.5 mL min^-1^ with a gradient from deionised water with 0.1% formic acid (FA) (eluent A) to acetonitrile with 0.1% FA (eluent B), from 2 %B to 30% B within 20 min, an increase to 75% B within 9 min, followed by a column cleaning-equilibration procedure. The QTOF settings were: negative electrospray ionisation, centroid mode, spectra rate 5 Hz, *m*/*z* range 50-1,300, end plate offset 500 V, capillary voltage 3,000 V, nebuliser (N_2_) pressure 3 bar, dry gas (N_2_) 12 L min^-1^ at 275°C, *m*/*z* 90 as low mass. In the MS mode, the quadrupole ion energy and collision energy were set to 4 eV and 7 eV, respectively. The AutoMSMS mode was used for MS/MS fragmentation, with *m*/*z*-dependent ramping of isolation widths and collision energies and N_2_ as collision gas. Sodium formate solution was introduced into the QTOF prior to each sample for *m*/*z* recalibration. Nine blanks were measured in addition. MetaboScape 2021b (Bruker Daltonics) was used for *m*/*z* recalibration and picking of metabolic features (each described by a RT and *m*/*z*; minimum peak height 750, at least 15 data points) in MS mode, applying the T-ReX 3D algorithm. Features probably belonging to the same metabolite, i.e., [M–H]^-^, [M+Cl]^-^, [M+HCOOH–H]^-^, [2M–H]^-^ and [M–H–H_2_O]^-^ions as well as their isotopes and charge states, were sorted together in so-called buckets, using 0.8 as correlation coefficient threshold. For each bucket, the feature with the highest intensity was used for quantification. Peak heights were divided by those of mefenamic acid ([M-H]^-^), means of blanks subtracted and data divided by the sample dry mass. Features (co-)eluting at RT < 1.25 min, i.e., close to or in the injection peak, including mostly primary metabolites, were omitted. For identification, those features, for which MS/MS spectra were available, were searched in a laboratory-internal library, comparing RTs, MS and MS/MS spectra. Additionally, some *C*-/*O*-glycosidic flavonoids, for which no reference standards were available, were putatively identified based on published fragmentation patterns (Ferreres et al. 2007; Waridel et al. 2001), specifically searching for those ones that have been previously found in wheat flag leaves (Stallmann et al. 2020).

### 2.7 Statistical Analyses

Depending on the measured trait, the final sample sizes were reduced to *n*=8 or *n*=9 in some groups due to mistakes during harvest and photosynthesis measurement, technical issues during chemical analyses and a malformed leaf for which no photosynthesis could be measured. Statistical analyses were done in R 4.4.2 (R Core Team 2024). The significance threshold was set to α = 0.05. Box-whisker plots were created using the “boxplot” function of the R package *graphics*. Measured traits were compared between the treatment groups within the two time points, applying (generalised) linear models (LMs or GLMs; R package *glmmTMB*) or weighted least squares (WLS) models with inverse group variances as weights (“lm”, R package *stats*). All models were fitted with the fixed factors watering (factor levels at T1: C, D; at T2: C-C, C-WL, D-WL) and fertilisation (factor levels: L, N) as well as their interaction term. For model diagnostics, the R packages *DHARMa* and *performance* were used; *p*-values were calculated based on type II tests (“Anova”, R package *car*), i.e., Wald *Χ²*-tests for (G)LMs and *F*-tests for WLS models. In case of (marginally) significant factors or interactions, manual contrasts with Bonferroni-Holm adjustment (R package *emmeans*) were applied, comparing the two fertilisation levels within the water control group (T1: C; T2: C-C) as well as contrasting each drought/ waterlogging group to the control group within fertilisation treatments.

Non-metric multidimensional scaling (NMDS) analyses were performed to compare the metabolite profiles of primary metabolites (LC-FLD, GC-MS) and the metabolic fingerprints (LC-MS) between samples, using Wisconsin double standardisation of square root-transformed data and Kulczynski distances (R package *vegan*). For each analytical platform, an overall NMDS analysis (all samples) as well as time point-specific analyses were done.

To trace responses of individual metabolites/ features to fertilisation and the watering treatments, concentrations were compared between the normal fertilisation and low fertilisation group within time points as well as between the drought/ waterlogging treatments and corresponding well-watered controls within time points and fertilisation regimes using Mann-Whitney *U*-tests followed by Benjamini-Hochberg correction of *p*-values (functions wilcox.test and p.adjust, R package *stats*) (Benjamini and Hochberg 1995). Mean fold changes for the fertilisation effects were computed as the mean concentration in the normal fertilisation group divided by the mean concentration in the low fertilisation group for each time point; for the fold changes related to the effects of the watering treatments, mean concentrations of the drought/ waterlogging treatment group were divided by the mean concentration in the corresponding control (T1: C, T2: C-C). For the fold changes, features that did not occur in both groups were excluded. Metabolites/ features were considered as being quantitatively modulated by the fertilisation or watering treatment if their concentrations significantly differed between groups (based on unadjusted *p*-values) and fold changes were < 0.67 (< −0.58 on log_2_ scale) or > 1.5 (log_2_: > 0.58); *p*-values and fold changes were visualised as volcano plots (Cui and Churchill 2003). Qualitative modulation was defined as occurrence in only one group in at least half of the replicates. The fold changes of those metabolites/ features, which were quantitatively modulated by the watering treatment in at least one pairwise comparison, were subjected to average linkage hierarchical clustering with Euclidean distances, clustering over group comparisons and metabolites/ features with the software Cluster 3.0 (de Hoon et al. 2004), followed by visualisation as cluster heatmap with Java Treeview (Saldanha 2004). For (partially) identified metabolites, fold changes of the watering treatment were plotted on a metabolic map (Schweiger et al. 2014; Schweiger et al. 2023) based on information from KEGG pathways (Kanehisa and Goto 2000) and additional information on biosynthesis of benzoxazinoids (Wouters et al. 2016).

## 3 Results

### 3.1 Effects of Watering and Fertilisation on Biomass

At the end of the drought phase (i.e., time point T1), the interaction between watering and fertilisation had a significant effect on the shoot dry biomass and a marginally significant effect on the root dry biomass (Table 1). In well-watered plants, the shoot biomass was higher under normal than under low fertilisation, while the root biomass did not differ between these groups (Figure 2A,B). Both shoot and root biomass were lower under drought compared to well-watered conditions, especially under normal fertilisation (only marginally significant at low fertilisation). The shoot-to-root ratio was significantly higher in normally than in low fertilised plants, but not affected by watering or the interaction (Table 1, Figure 2C).

**FIGURE 2.**
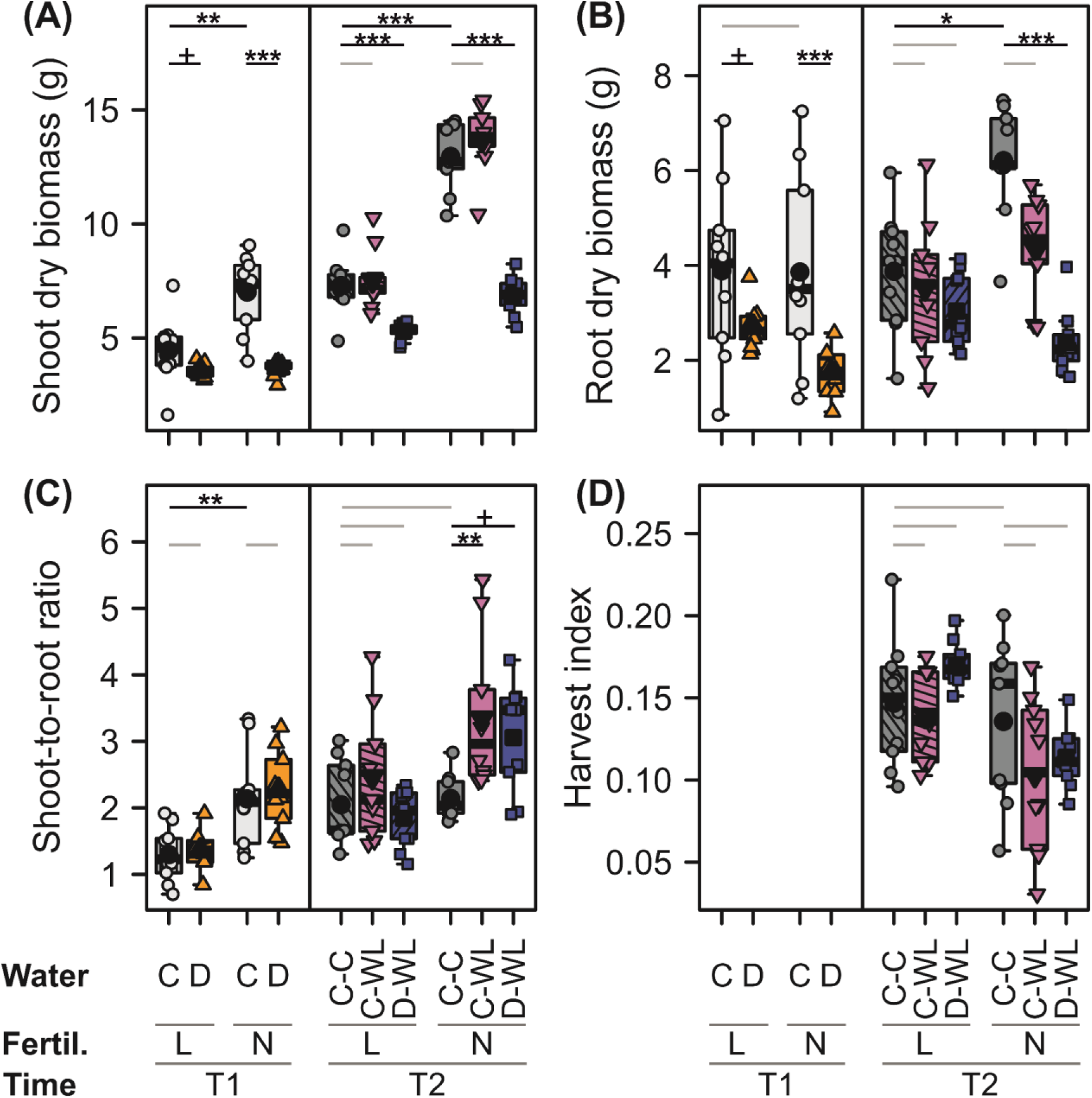
Shoot (A) and root (B) dry biomass, (C) shoot-to-root ratio and (D) harvest index (i.e., ratio of inflorescence dry biomass to total shoot dry biomass) of *Triticum aestivum*, calculated per pot with two plants each. Plants were subjected to different watering regimes (well-watered control, C; drought, D) in the drought phase, while in the subsequent waterlogging phase they were well-watered (C) or waterlogged (WL). Two fertilisation treatments were applied (low, N; normal, N). Plants were harvested at two time points (end of drought phase, T1; end of waterlogging phase, T2). The harvest index was only calculated for T2, at which ears had emerged. Data are given as box-whisker plots, with raw data as small symbols, medians as thick horizontal lines, means as large black symbols, interquartile ranges (IQR) as boxes (hatched for the L treatment) and whiskers extending to the maximum data points within 1.5 x IQR. Results of statistical tests are given in Table 1; if factors were (marginally) significant, manual contrasts are shown as horizontal lines [grey: not significant; black: (marginally) significant] with significance levels above the lines: + *p* < 0.1, * *p* < 0.05, ** *p* < 0.01, *** *p* < 0.001]; *n*=9-10.

**TABLE 1.**
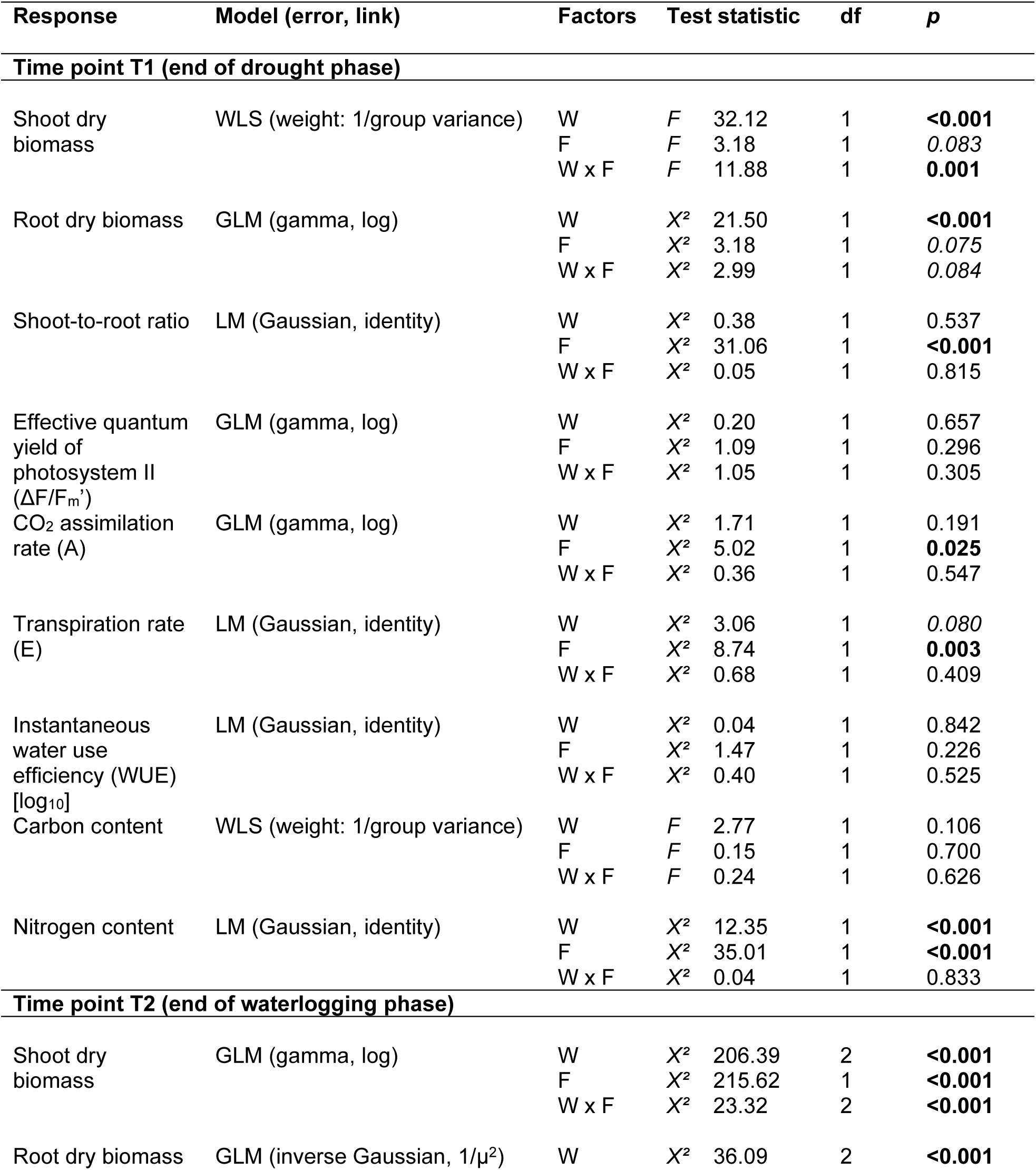
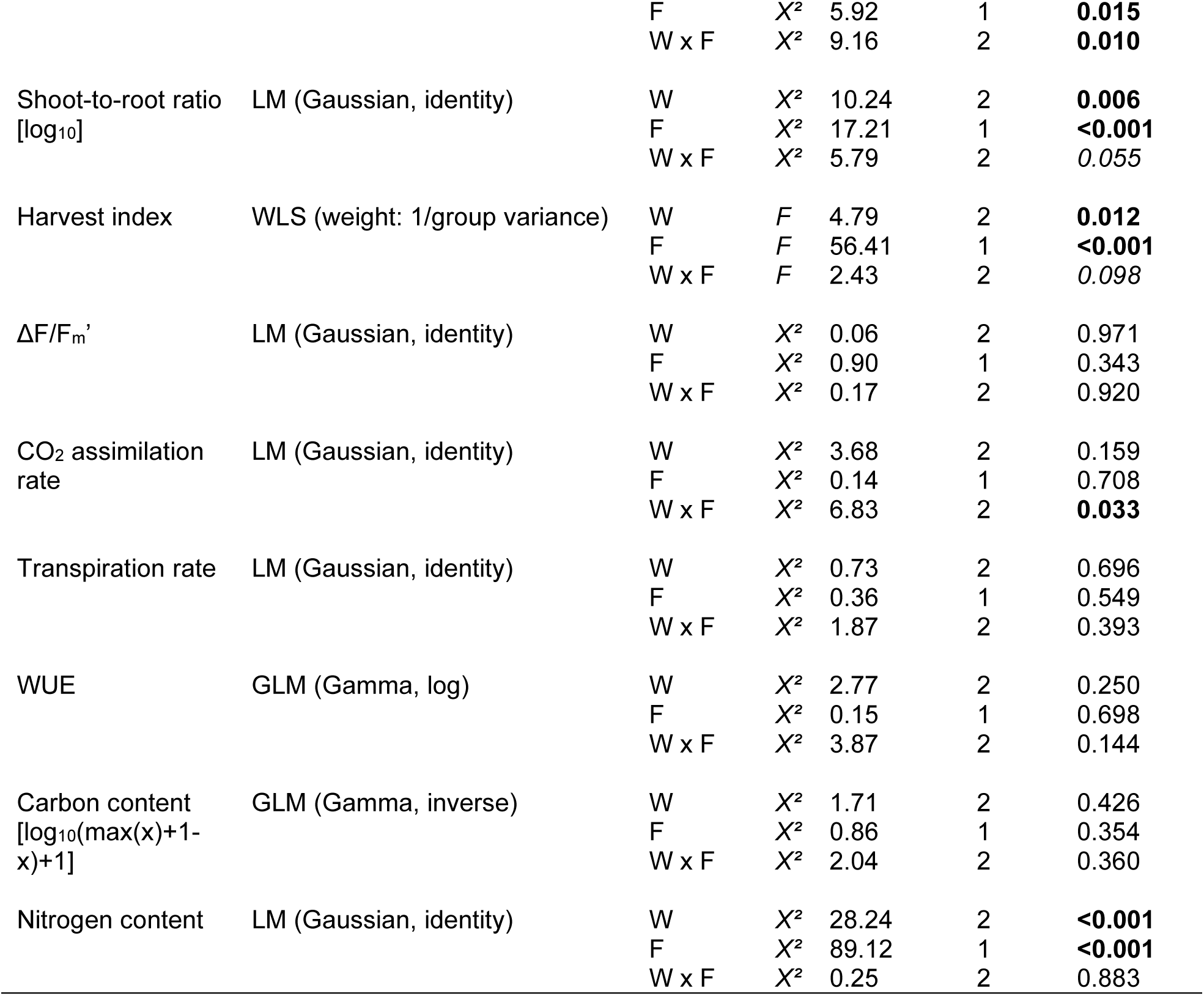
Effects of the factors watering (W) and fertilisation (F) as well as their interaction (WxF) on traits of *Triticum aestivum* (with data transformation, if applicable, in brackets). The model type (LM: linear model; GLM: generalised linear model; WLS: weighted least squares model), error family and link function (for LMs, GLMs), type of weights (for WLS) as well as the statistical results (*Χ²* or *F* values, df, *p*-values) are given, with marginally significant *p*-values (< 0.1) in italics and significant (< 0.05) ones in bold.

At the end of the waterlogging phase (T2), both shoot and root dry biomass were significantly affected by the watering x fertilisation interaction (Table 1). In well-watered plants (C-C), both traits were higher under normal than under low fertilisation (Figure 2A,B). Plants subjected to the drought-waterlogging sequence (D-WL) had a lower shoot biomass than C-C plants, while shoots of plants subjected to waterlogging without prior drought (C-WL) were similar to C-C plants (Figure 2A). Similarly, for the root biomass, a reduction was found in D-WL compared to C-C plants under normal fertilisation, while no differences were found at low fertilisation (Figure 2B). The watering x fertilisation interaction also affected (marginal significance) the shoot-to-root ratio and harvest index at T2, with the shoot-to-root ratio being higher in the C-WL (significant) and D-WL (marginally significant) group than in the C-C group, but only under normal fertilisation (Table 1, Figure 2C,D).

### 3.2 Effects of Watering and Fertilisation on Photosynthesis

At T1, ΔF/F_m_’ was neither affected by watering, fertilisation nor their interaction (Table 1, Figure 3A). There was a marginally significant effect of watering on the transpiration rate, while fertilisation showed significant impacts on the CO_2_ assimilation and the transpiration rate (Table 1); however, the post hoc manual contrasts were not significant for both traits (Table 1, Figure 3B,C). The WUE was not affected by any of the factors (Table 1, Figure 3D). At T2, there was a significant watering x fertilisation interaction effect on the CO_2_ assimilation rate, without significant post hoc differences, while neither ΔF/F_m_’, the transpiration rate nor the WUE were influenced by any of the factors (Table 1, Figure 3A-D).

**FIGURE 3.**
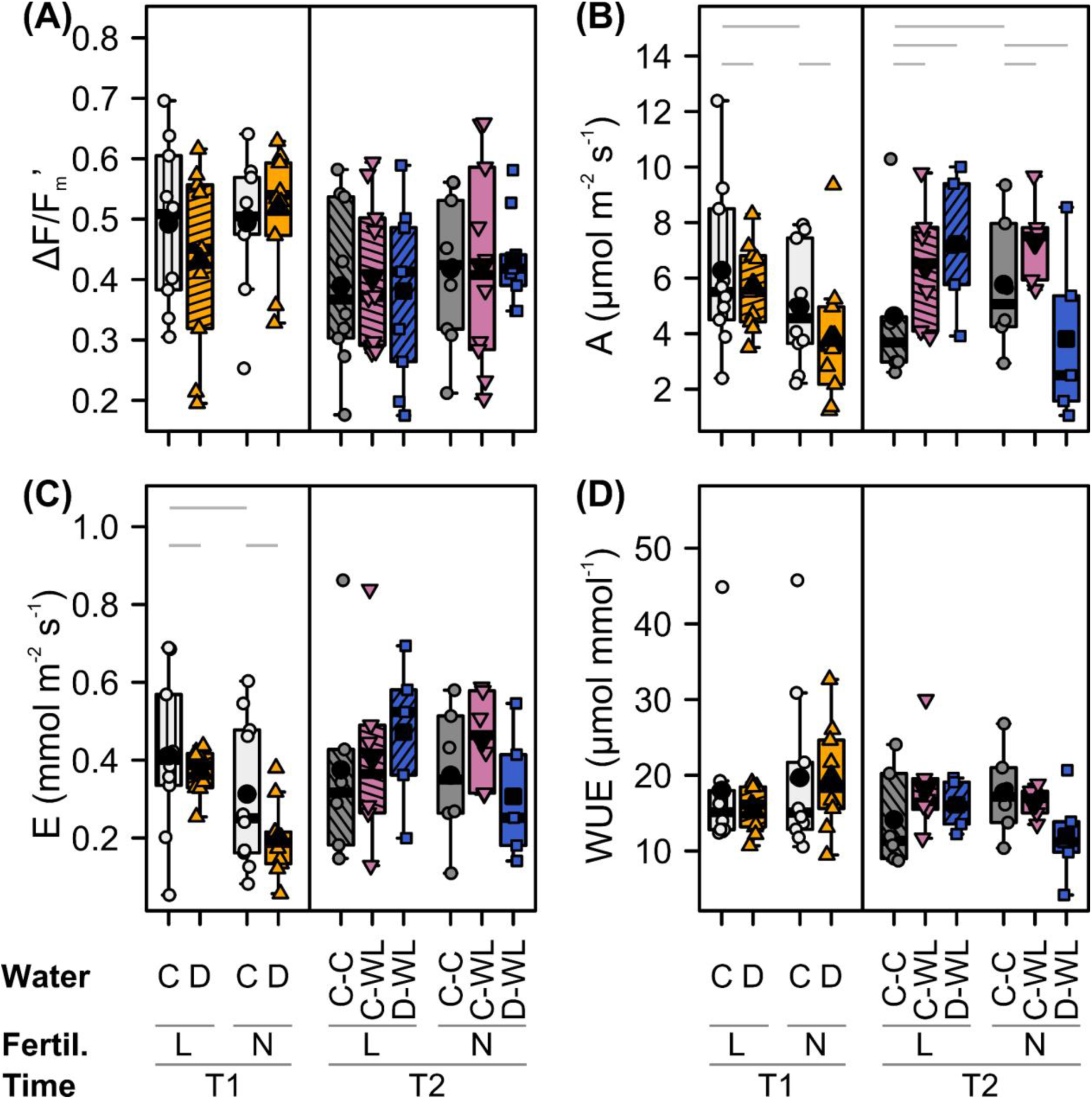
Effective quantum yield of photosystem II (ΔF/F_m_’; A), CO_2_ assimilation rate A (B), transpiration rate E (C) and instantaneous water use efficiency WUE (D) of *Triticum aestivum* leaves. Plants were subjected to different watering regimes (well-watered control, C; drought, D) in the drought phase, while in the subsequent waterlogging phase they were well-watered (C) or waterlogged (WL). Two fertilisation treatments were applied (low, N; normal, N). Measurements were done at two time points (end of drought phase, T1; end of waterlogging phase, T2). Data are given as box-whisker plots, with raw data as small symbols, medians as thick horizontal lines, means as large black symbols, interquartile ranges (IQR) as boxes (hatched for the L treatment) and whiskers extending to the maximum data points within 1.5 x IQR. Results of statistical tests are given in Table 1; if factors were (marginally) significant, manual contrasts are shown as horizontal lines [grey: not significant]; *n*=9-10 for ΔF/F_m_’; *n*=5-10 for the other traits.

### 3.4 Effects of Watering and Fertilisation on Leaf Chemistry

At both time points, the foliar carbon content was not affected by watering, fertilisation or their interaction. In contrast, watering and fertilisation had significant impacts on the nitrogen content (Table 1, Figure 4). In well-watered plants, it was significantly higher under normal than under low fertilisation at both time points (Figure 4B). Moreover, plants subjected to drought (T1) and to the drought-waterlogging sequence, but not those exposed to waterlogging only (T2), showed higher nitrogen contents than well-watered plants.

**FIGURE 4.**
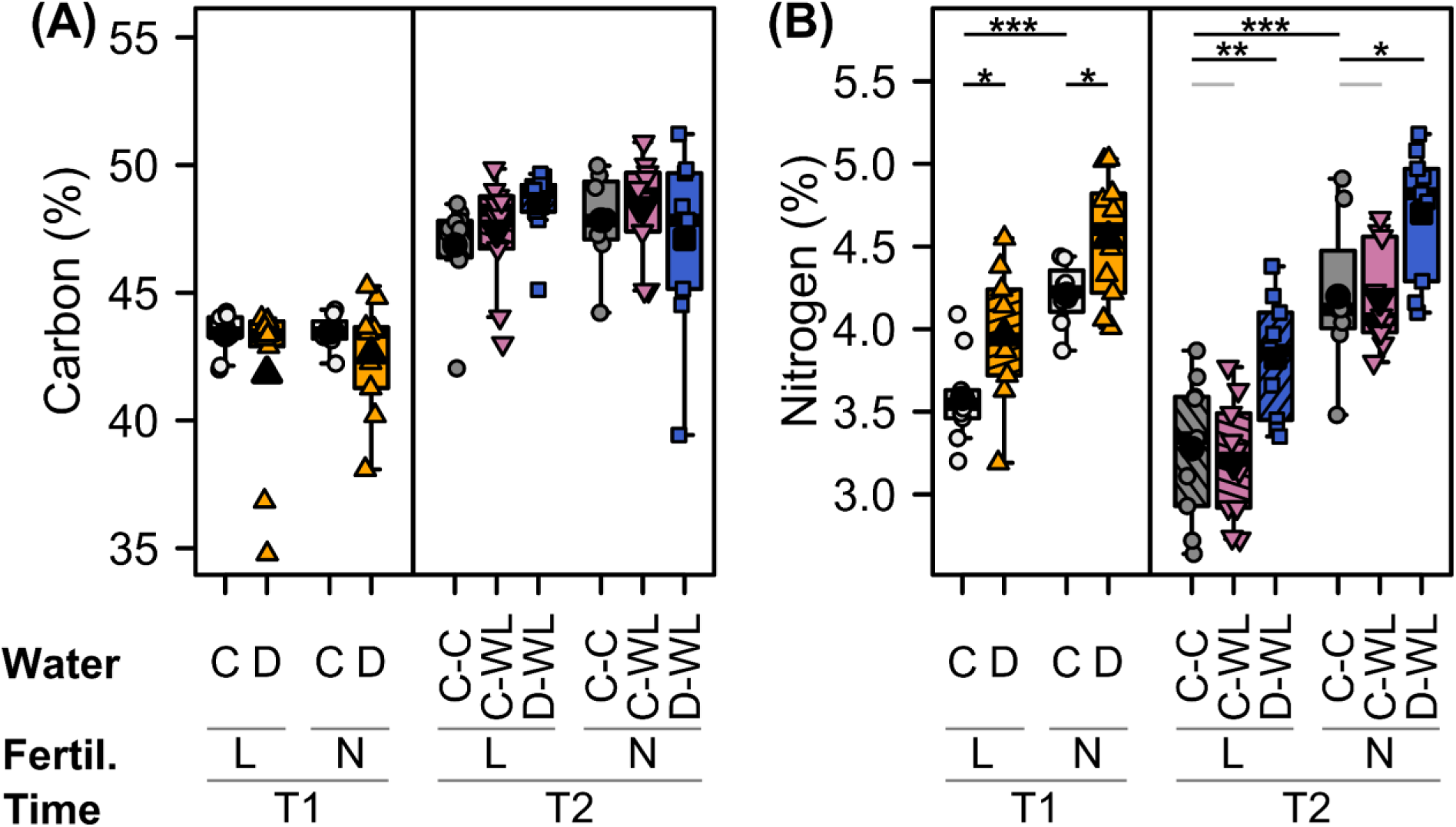
Carbon (A) and nitrogen (B) contents of *Triticum aestivum* leaves. Plants were subjected to different watering regimes (well-watered control, C; drought, D) in the drought phase, while in the subsequent waterlogging phase they were well-watered (C) or waterlogged (WL). Two fertilisation treatments were applied (low, N; normal, N). Leaves were harvested at two time points (end of drought phase, T1; end of waterlogging phase, T2). Data are given as box-whisker plots, with raw data as small symbols, medians as thick horizontal lines, means as large black symbols, interquartile ranges (IQR) as boxes (hatched for the L treatment) and whiskers extending to the maximum data points within 1.5 x IQR. Results of statistical tests are given in Table 1; if factors were (marginally) significant, manual contrasts are shown as horizontal lines [grey: not significant; black: (marginally) significant] with significances above the lines: * *p* < 0.05, *** *p* < 0.001]; *n*=8-10.

Profiling of primary metabolites in the leaves revealed 22 amino acids (21 primary, one secondary), 10 further primary metabolites (carbohydrates, organic acids, *myo*-inositol) and one inorganic acid (Supporting Information S1: Table S1). In addition, via metabolic fingerprinting we found 2,372 metabolic features (1,725 features at T1; 1,963 at T2), of which 14 could be identified, comprising two benzoxazinoids and twelve flavonoids.

The metabolite patterns, assessed per analytical platform via non-metric multidimensional scaling (NMDS), largely overlapped between the groups, but also pointed to some differences between fertilisation and water treatment groups within time points (Figure S1). In well-watered plants, the concentrations of many metabolites/ features were affected by fertilisation (Figure S2). Specifically, various flavonoids had lower concentrations in plants under normal fertilisation compared to plants under low fertilisation at both time points, while benzoxazinoids (at T1) and many amino acids (at T2) were accumulated at normal fertilisation. Moreover, various metabolites/ features were affected by drought, waterlogging and/ or the drought-waterlogging sequence, with some showing lower and others higher concentrations than in the well-watered controls (Figures 5, S3). Several primary metabolites and unknown metabolic features, but neither benzoxazinoids nor flavonoids, were modulated by one or more of the watering treatments (Figures 5, 6). At the end of the drought phase (T1), asparagine, glutamine and proline showed higher concentrations, while *myo*-inositol exhibited lower concentrations under drought independent of fertilisation (Figures 5, 6). Some further metabolites responded only under low or normal fertilisation. At the end of the waterlogging phase (T2), less metabolites were modulated by waterlogging if plants were well-watered before (C-WL) compared to previously drought-exposed (D-WL) plants (Figure 5). Several amino acids had lower concentrations in the C-WL than in the C-C group, including asparagine in both fertilisation groups and proline, serine and citrulline under normal fertilisation. The response to the drought-waterlogging sequence (D-WL versus C-C) strongly depended on fertilisation. While several amino acids showed higher concentrations after drought-waterlogging in the low fertilisation group, no amino acids but phosphoric acid and malic acid as well as fructose showed lower concentrations in the normal fertilisation group. Mapping the fold changes of identified metabolites on pathways revealed that several biosynthetically related metabolites responded similarly to the water treatments (Figure 6). The most pronounced responses appeared in a cluster of amino acids (asparagine, glutamine, GABA, proline), showing higher concentrations at the end of the drought phase, while concentrations were partly higher or lower at the end of the waterlogging phase. Several other amino acids (e.g., tryptophan, tyrosine, homoserine, leucine) only responded with higher concentrations to the drought-waterlogging sequence in the low fertilisation group. In and close to the glycolysis pathway, *myo*-inositol showed lower concentrations in drought-exposed plants of both fertilisation regimes at T1, while glucose and fructose responded with higher concentrations in drought-exposed plants at T1, especially under normal fertilisation, and lower concentrations in drought-waterlogging-exposed plants of normal fertilisation at T2 (Figure 6). Overall, clustering of fold changes revealed that metabolic responses to waterlogging and to the drought-waterlogging sequence under low fertilisation at T2 were most similar (Figure S3). Likewise, responses to drought at T1 under low fertilisation were similar to responses to drought followed by waterlogging (T2) under normal fertilisation. Metabolic response to drought at T1 as well as to waterlogging at T2, both under normal fertilisation, were most distinct to the other group comparisons. Within watering treatments, the low and normal fertilisation groups showed quite different overall metabolic responses, as they did not cluster together, neither for the response to the single watering regimes nor the drought-waterlogging sequence (Figure S3).

**FIGURE 5.**
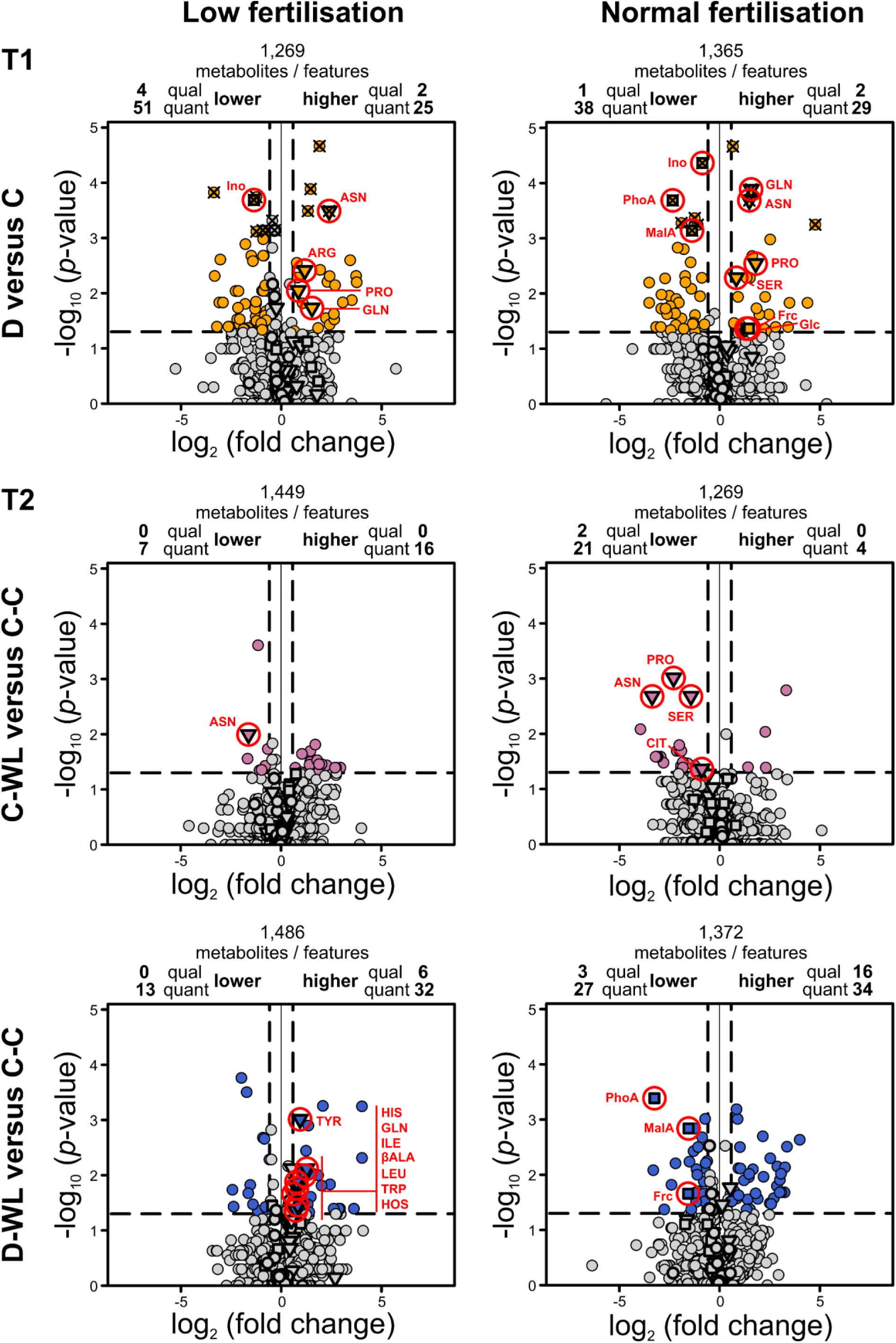
Modulation of metabolites and metabolic features in *Triticum aestivum* leaves by drought and/ or waterlogging, shown as volcano plots. Plants were subjected to different watering regimes (well-watered control, C; drought, D) in the drought phase, while in the subsequent waterlogging phase they were well-watered (C) or waterlogged (WL). Two fertilisation treatments were applied (low, N, first column; normal, N, second column). Leaves were harvested at two time points (end of drought phase, T1, first row; end of waterlogging phase, T2, second and third row). For each group comparison, *p*-values (Mann-Whitney *U*-tests, -log_10_) and mean fold changes (log_2_) of metabolites/ features occurring in both groups are plotted. Coloured symbols represent modulated metabolites/ features outside the dashed cutoff lines (horizontal: corresponding to *p*-value of 0.05; vertical: corresponding to fold changes of 0.67 and 1.5), with those on the left being lower and those on the right being higher in the treatment than in the control group. Non-modulated metabolites/ features are shown in grey. Symbol types indicate analytical platforms (triangles: LC-FLD, amino acids; squares: GC-MS, further primary metabolites; circles: LC-MS, mainly specialised metabolites). (Partially) identified metabolites are surrounded by thicker lines and labeled in red if they were quantitatively modulated (full names: Table S1); there were no identified metabolites among the qualitatively modulated features. Metabolites/ features that were significant after Benjamini-Hochberg correction of *p*-values are indicated with crosses. The numbers of qualitatively and quantitatively modulated features are given at the top. For one comparison (T2, L, C-WL versus C-C), a non-significant outlier feature with a log_2_ fold change < −11 was omitted; based on *n*=9-10.

**FIGURE 6.**
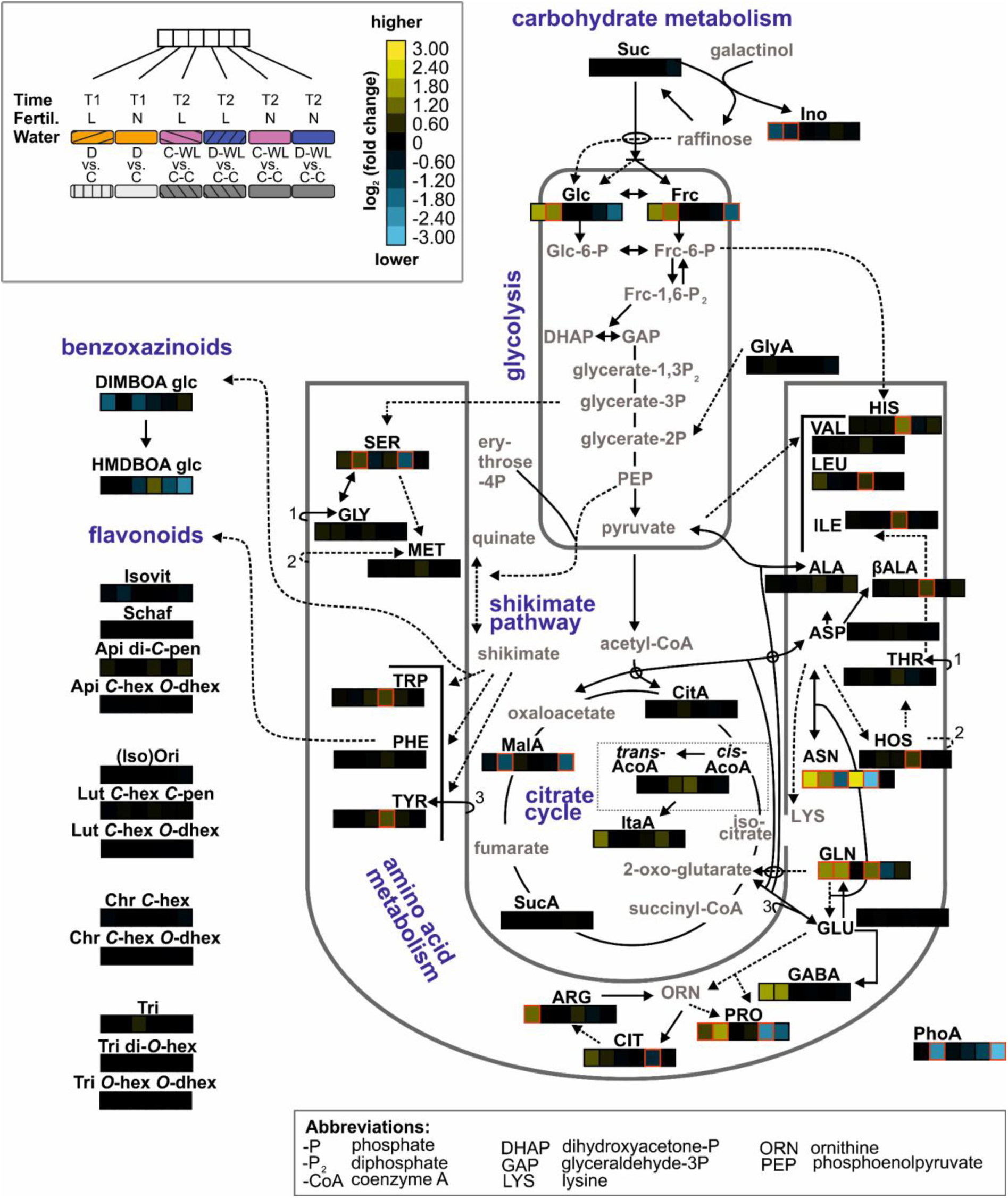
Mean fold changes (log_2_) of metabolites and metabolic features in leaves of *Triticum aestivum* plants that were subjected to drought and/ or waterlogging, displayed on a metabolic map. Plants were subjected to different watering regimes (well-watered control, C; drought, D) in the drought phase, while in the subsequent waterlogging phase they were well-watered (C) or waterlogged (WL). Two fertilisation treatments were applied (low, N; normal, N). Leaves were harvested at two time points (end of drought phase, T1; end of waterlogging phase, T2). Yellow and blue indicate higher and lower concentrations in the drought/ waterlogging compared to the control group, respectively. Metabolites, which were quantitatively modulated (*p*-value < 0.05 in Mann-Whitney *U*-test; log_2_-scaled fold changes < 0.58 or > 0.58) are highlighted with red squares. Some intermediates are not shown (dashed arrows). For full names, see Table 1 (metabolites found in our study) and legend at the bottom (further metabolites); based on *n*=9-10.

## 4 Discussion

### 4.1 Effects of fertilisation

In our study, several traits of well-watered wheat plants were affected by fertilisation. As expected, the shoot and (at T2) root biomass and shoot-to-root ratio (at T1) were higher under normal than under low fertilisation, as also found in various other crop species under enhanced fertilisation (Lopez et al. 2023). The fact that normally fertilised plants use fertiliser for their shoot growth but produce relatively less root biomass may be due to the lower effort needed to acquire nutrients at higher fertilisation and/ or be related to differences in plant size affecting biomass allocation (Poorter et al. 2012). Contrary to our expectation, but consistent with a study on durum wheat (*Triticum turgidum* ssp. *durum*) (Cabrera-Bosquet et al. 2009), the harvest index was not affected by fertilisation. Moreover, although we had assumed positive effects of higher fertilisation on CO_2_ assimilation – in line with findings for durum and bread wheat (Cabrera-Bosquet et al. 2009; Guo et al. 2019) – the absence of such effects in our study may indicate that photosynthesis was not nutrient-limited. As hypothesised, foliar N contents were higher in normally fertilised plants, suggesting that N was taken up and transported to the leaves, while growth-mediated dilution did not prevent foliar N accumulation under these conditions.

Along with biomass allocation shifts, we found various metabolic differences between leaves from plants of the normal versus the low fertilisation group. Nutrients are important for the photosynthetic machinery, deliver building blocks for metabolites and enzymes, are part of the energy metabolism and can modulate regulatory networks affecting resource allocation in metabolic pathways (Tariq et al. 2023). When various nutrients differ in the provided fertiliser amount, as in our study, the metabolic outcome probably results from a complex, multidimensional interplay of nutrients, growth and metabolism. We found higher concentrations of various amino acids (at T2) and lower concentrations of different flavonoids under normal compared to low fertilisation, similarly as previously reported (Heyneke et al. 2017; Zhang et al. 2017). Under nutrient depletion, the flavonoid pathway may be upregulated, as found in *Arabidopsis thaliana* (Brassicaceae) (Lillo et al. 2008).

### 4.2 Role of fertilisation in plant responses to drought

Drought had various effects on wheat. As expected, the shoot and root dry biomass were lower under drought, as likewise reported in numerous other studies of wheat (Wang et al. 2017; Xu et al. 2025). This response indicates that plant functions necessary for growth are constrained under water limitation. However, the shoot-to-root ratio was not affected by drought in our present study, although plants often invest more in root than in shoot biomass under drought, likely to capture water more effectively (Eziz et al. 2017; Poorter et al. 2012). Moreover, contrary to our hypothesis, neither photosynthesis nor the WUE were affected by drought, indicating that the photosynthetic apparatus was still functional and stomata were open under drought. While photosynthesis is often reduced under drought (Chaves et al. 2003; Flexas and Medrano 2002; Wang et al. 2017), this physiological response may depend on the drought severity. Our plants may have responded to the drought conditions by investing less in new leaves, while maintaining photosynthesis in existing leaves. Differences in leaf aging and sink-source transition in younger leaves of well-watered versus drought-exposed plants (Chaves et al. 2003) may also have contributed to the high CO_2_ assimilation in both treatment groups. At more severe drought, stomatal and non-stomatal factors may restrict photosynthesis.

As expected, foliar N contents were higher under drought, in line with a previous finding reporting higher N contents in wheat shoots (Pons and Müller 2022). While the uptake of water and N was probably hampered under water limitation, a slower growth may have (over)compensated this, leading to less N dilution and thus higher N contents. As the N demand depends on plant size, N contents may be higher if plants are smaller under water deficit (Gonzalez-Dugo et al. 2010). Moreover, N-containing metabolites are often accumulated under drought (Itam et al. 2020; Wang et al. 2017). In accordance, we found higher concentrations of several amino acids, including proline, in leaves of drought-exposed plants. Proline, other amino acids and further small metabolites accumulate under challenges such as drought, act as osmolytes and protect cells (Chaves et al. 2003; Fàbregas and Fernie 2019; Szabados and Savouré 2010). The higher concentrations of asparagine found in our study under drought may be related to an earlier onset of senescence, as suggested previously (Itam et al. 2020). Benzoxazinoid and flavonoid concentrations were not affected by drought, while an accumulation of leaf benzoxazinoids and a change in flavonoids had been found under more severe, i.e., pulsed, drought in another study on wheat (Stallmann et al. 2020). Also, in leaves of wheat seedlings, benzoxazinoids metabolism was up-regulated after 18 days of drought (Hao et al. 2026). Taken together, the studies suggest that the timing and/ or intensity of drought are decisive.

The potential of nutrient application to help plants cope with drought stress, e.g., via improved antioxidant capacity and osmoregulation, has been recently highlighted (Drobnitch et al. 2025; Swain et al. 2023). However, in our experiment plants did not benefit from higher fertilisation when exposed to drought. In contrast, these plants showed a stronger reduction in shoot and root biomass in relation to well-watered plants compared to plants fertilised less. Lower nutrient availability may have exacerbated the lower nutrient uptake capacities under drought. Under well-watered conditions, the normally fertilised plants had a higher biomass and thus probably higher nutritional needs, which were likely harder to satisfy under drought. Similarly, normally fertilised plants may have more building blocks for drought-induced metabolic changes, but these responses may be diluted in larger plants. Such mechanisms may explain the overall comparable foliar metabolic responsiveness to drought in plants of the two fertilisation groups. Despite overlapping responses, several drought-induced metabolic changes differed, indicating that nutrition and plant growth together may have shaped flag leaf metabolism. Other studies on wheat revealed fertiliser-dependent effects of drought on various traits, with the highest N fertiliser application not necessarily maximising grain yield (Drobnitch et al. 2025; Ru et al. 2024; Ru et al. 2023).

### 4.3 Role of fertilisation in plant responses to waterlogging

Compared to drought, waterlogging had less effects on wheat and neither shoot or root dry biomass, CO_2_ assimilation rate nor foliar N content were affected. Other studies revealed various adverse effects of waterlogging on crops including wheat on these traits (Cid et al. 2024; Herzog et al. 2016; Tian et al. 2021). The low responsiveness of plants to waterlogging in our experiment may have been due to the relatively short waterlogging duration (i.e., 8 days). Indeed, the duration of waterlogging is decisive for plant responses (Tian et al. 2021). In line with our expectation, the shoot-to-root ratio was higher under waterlogging, at least under normal fertilisation, suggesting that root systems may have been partially destroyed. A lower root mass and higher shoot-to-root ratios are characteristic for waterlogged plants including wheat (Herzog et al. 2016; Poorter et al. 2012). Moreover, as expected, we found several metabolic changes in response to waterlogging. Some amino acids had lower concentrations in leaves of waterlogged compared to control plants. Likewise, most foliar amino acids were less concentrated after a waterlogging period of 12 days in wheat (Cid et al. 2024), but, for example, concentrations of glutamic acid and GABA were higher in seedlings waterlogged for 3 days (Komatsu et al. 2023). These findings indicate that not only the duration but also the timing of the waterlogging during development may be crucial for metabolic responses. Sugars and further organic acids were less affected in our study compared to a study where wheat was waterlogged 4 days longer (Cid et al. 2024). Due to the overall low responsiveness of wheat to waterlogging in our experiment, we cannot assess the extent to which fertilisation may influence the responses. In cotton (*Gossypium hirsutum*, Malvaceae), benefits of additional N fertilisation for waterlogged plants were revealed, which depended on N application rates and the waterlogging duration (Yang et al. 2026). In wheat, waterlogging tolerance has been linked with the capacity to take up and utilise sulphur (Hussain et al. 2022) and potassium attenuated negative effects of waterlogging (Gao et al. 2021). However, effects of waterlogging and alleviating effects of fertilisation in pot experiments may be limited, as nutrient loss under waterlogging and plant growth are restricted under such conditions (Xu et al. 2026).

### 4.4 Role of fertilisation in plant responses to drought followed by waterlogging

The effects of the drought-waterlogging sequence on wheat were similar to – and in some cases more severe than – the impacts of drought and/ or waterlogging alone, partially supporting our hypotheses. The pronounced reduction of shoot and root biomass in plants subjected to the sequence under normal fertilisation may have been due to drought-induced limitations of plant growth as well as (beginning) root decay under waterlogging. In line with our findings, lower shoot and root biomass under similar water regimes have been reported in wheat and in maize, while the harvest index was not affected in wheat (Keya et al. 2025; Schweiger et al. 2023). The lack of effects of the drought-waterlogging sequence on photosynthesis traits in our current and a previous study (Schweiger et al. 2023) indicates ongoing photosynthesis in the leaves. In contrast, longer durations of drought followed by waterlogging, i.e., 15 or 21 days each, impaired photosynthesis in wheat and maize, but not in barley (*Hordeum vulgare*, Poaceae) (Keya et al. 2025; Saha and Johnson 2025). The higher foliar N contents that we found in plants subjected to the drought-waterlogging sequence likely mainly resulted from the preceding drought phase, with less N dilution in smaller plants and a potential accumulation of N-containing metabolites. After longer drought-waterlogging periods (each 15 days), lower shoot N contents have been reported both for wheat and barley (Saha and Johnson 2025). Lower fructose concentrations in leaves of normally fertilised plants in response to our drought-waterlogging sequence partly fit to our previous finding (Schweiger et al. 2023). Proline concentrations were slightly lower after drought-waterlogging in the current study, but slightly higher in our earlier study with partly similar treatments (Schweiger et al. 2023) and much higher after drought-waterlogging (21 days each) in maize leaves (Keya et al. 2025). This discrepancy is again likely related to different stress intensities, while species-specific responses may also play a role.

Surprisingly, the impacts of drought-waterlogging on biomass and the shoot-to-root ratio were more pronounced in plants under normal fertilisation, while N contents were similarly affected in both fertilisation groups. However, metabolic responses were distinct. Although low fertilisation may have exacerbated nutrient deficiencies both in the drought and following waterlogging phase, plant size-dependent nutritional requirements and dilution of pool sizes of metabolic building blocks must probably be taken into account to understand these results, as discussed for fertilisation and drought above. For example, drought followed by waterlogging induced higher concentrations of amino acids only in leaves of plants under low fertilisation, which may indicate that N demands for accumulation of these metabolites were met in these smaller plants.

## 5 Conclusions

Taken together, our study revealed numerous effects of challenging watering availabilities on wheat. In addition to impacts of drought or waterlogging alone, we uncovered effects of drought and subsequent waterlogging, which is highly relevant to understand consequences of ongoing climate change for (agricultural) plants. Discrepancies between studies, even with the same or closely related plant species, are probably largely due to different stress intensities, highlighting the need for more studies incorporating such factors, e.g., different durations of watering treatments within one study. Moreover, our experiment and other studies revealed that fertilisation modulates plant responses to drought and waterlogging, at least in part. When evaluating interactions between nutrient and water availability, plant nutritional demands have to be considered. While supplemental fertilisation can improve plant performance and is considered in agricultural production of crops suffering from drought and waterlogging (Drobnitch et al. 2025; Xu et al. 2026), it also raises concerns about environmental pollution (Penuelas et al. 2023). Thus, agronomic, economic and environmental aspects as well as nutrient demand and supply have to be balanced in agricultural management (Gonzalez-Dugo et al. 2010). Ideally, precision fertilisation strategies, considering type and concentration of nutrients as well as timing of application, should be embedded in soil-nutrient-climate frameworks (Drobnitch et al. 2025; Lee et al. 2026). Our study contributes to a deeper understanding of the interplay between water availability and fertilisation-based nutrition in wheat.

## Acknowledgements

We thank Viktorija Bolelaja, Pia Deimann, Stella Lohoff, Leona Meyer, Sabrina Nufer, Kristina Runte and Lina Solmecke for practical help in the experiments, Karin Djendouci for help with the elementar analyses as well as Caroline Pons for insights from pre-tests and fruitful discussions.

## Data Availability Statement

The raw LC-MS data will be made available on MetaboLights (Yurekten et al. 2024). All other raw data are part of the Supporting Information of this publication.

## Supporting Information

Additional supporting information can be found online in the Supporting Information section.

## References

Arif, M., S. Fiaz, W. M. W. W. Kandegama, S. Sheth, and L. Li. 2025. “Evaluating the Impacts of Environmental Stresses on Agriculture in the Context of Climate Resilience.” Plant Molecular Biology 115, no. 4: 73.

Benjamini, Y., and Y. Hochberg. 1995. “Controlling the False Discovery Rate: A Practical and Powerful Approach to Multiple Testing.” Journal of the Royal Statistical Society B 57, no. 1: 289–300.

Cabrera-Bosquet, L., R. Albrizio, J. L. Araus, and S. Nogués. 2009. “Photosynthetic Capacity of Field-grown Durum Wheat Under Different N Availabilities: A Comparative Study From Leaf to Canopy.” Environmental and Experimental Botany 67, no. 1: 145–152.

Chaves, M. M., J. P. Maroco, and J. S. Pereira. 2003. “Understanding Plant Responses to Drought – From Genes to the Whole Plant.” Functional Plant Biology 30, no. 3: 239–264.

Cid, G. A., D. Francioli, S. Kolb, Y. A. Tandron Moya, N. von Wirén, and M.-R. Hajirezaei. 2024. “Transcriptomic and Metabolomic Approaches Elucidate the Systemic Response of Wheat Plants Under Waterlogging.” Journal of Experimental Botany 75, no. 5: 1510–1529.

Cui, X., and G. A. Churchill. 2003. “Statistical Tests for Differential Expression in cDNA Microarray Experiments.” Genome Biology 4: 210.

de Hoon, M. J. L., S. Imoto, J. Nolan, and S. Miyano. 2004. “Open Source Clustering Software.” Bioinformatics 20, no. 9: 1453–1454.

Dietz, K.-J., C. Zörb, and C.-M. Geilfus. 2021. “Drought and Crop Yield.” Plant Biology 23, no. 6: 881–893.

Drobnitch, S. T., T. C. Donovan, J. A. Wenz, N. E. Flynn, M. E. Schipanski, and L. H. Comas. 2025. “Can Nitrogen Availability Impact Plant Performance Under Water Stress? A Review of Traits, Mechanisms, and Whole Plant Effects.” Plant and Soil 511, no. 1–2: 45–67.

European Commission. 2020. “A Farm to Fork Strategy for a Fair, Healthy and Environmentally-friendly Food System.” https://eur-lex.europa.eu/legal-content/EN/TXT/?uri=celex%3A52020DC0381.

Eziz, A., Z. Yan, D. Tian, W. Han, Z. Tang, and J. Fang. 2017. “Drought Effect on Plant Biomass Allocation: A Meta-analysis.” Ecology and Evolution 7, no. 24: 11002–11010.

Fàbregas, N., and A. R. Fernie. 2019. “The Metabolic Response to Drought.” Journal of Experimental Botany 70, no. 4: 1077–1085.

Farooq, M., A. Wahid, N. Kobayashi, D. Fujita, and S. M. A. Basra. 2009. “Plant Drought Stress: Effects, Mechanisms and Management.” Agronomy for Sustainable Development 29, no. 1: 185–212.

Ferreres, F., A. Gil-Izquierdo, P. B. Andrade, P. Valentão, and F. A. Tomás-Barberán. 2007. “Characterization of *C*-glycosyl Flavones *O*-glycosylated by Liquid Chromatography–Tandem Mass Spectrometry.” Journal of Chromatography A 1161, no. 1–2: 214–223.

Flexas, J., and H. Medrano. 2002. “Drought-inhibition of Photosynthesis in C_3_ Plants: Stomatal and Non-stomatal Limitations Revisited.” Annals of Botany 89, no. 2: 183–189.

Gao, J., Y. Su, M. Yu, Y. Huang, F. Wang, and A. Shen. 2021. “Potassium Alleviates Post-anthesis Photosynthetic Reductions in Winter Wheat Caused by Waterlogging at the Stem Elongation Stage.” Frontiers in Plant Science 11: 607475.

Geng, R., M. Xu, L. Xu, G. Yan, and G. Cai. 2026. “Biological Mechanisms of Waterlogging Tolerance in Plants.” Plant, Cell & Environment 49, no. 2: 685–699.

Gonzalez-Dugo, V., J.-L. Durand, and F. Gastal. 2010. “Water Deficit and Nitrogen Nutrition of Crops. A Review.” Agronomy for Sustainable Development 30, no. 3: 529–544.

Guo, J., Y. Jia, H. Chen, L. Zhang, J. Yang, J. Zhang, X. Hu, X. Ye, Y. Li, and Y. Zhou. 2019. “Growth, Photosynthesis, and Nutrient Uptake in Wheat are Affected by Differences in Nitrogen Levels and Forms and Potassium Supply.” Scientific Reports 9, no. 1: 1248.

Hao, L. K., Z. Batyrshina, Y. Goldstein, A. Mishra, T. G. Köllner, B. Yaakov, and V. Tzin. 2026. “Dual Role of Benzoxazinoids in Plant Response to Combined Drought Stress and Aphid Feeding.” Plant Biology 28, no. 1: 45–57.

Herzog, M., G. G. Striker, T. D. Colmer, and O. Pedersen. 2016. “Mechanisms of Waterlogging Tolerance in Wheat – A Review of Root and Shoot Physiology.” Plant, Cell & Environment 39, no. 5: 1068–1086.

Heyneke, E., M. Watanabe, A. Erban, G. Duan, P. Buchner, D. Walther, J. Kopka, M. J. Hawkesford, and R. Hoefgen. 2017. “Characterization of the Wheat Leaf Metabolome During Grain Filling and Under Varied N-supply.” Frontiers in Plant Science 8: 2048.

Hummel, J., N. Strehmel, J. Selbig, D. Walther, and J. Kopka. 2010. “Decision Tree Supported Substructure Prediction of Metabolites From GC-MS Profiles.” Metabolomics 6, no. 2: 322–333.

Hussain, M. A., A. Naeem, S. Sulieman, B. Pitann, and K. H. Mühling. 2022. “Sulfur Uptake and Distribution, Grain Yield, and Quality of Hybrid and Inbred Winter Wheat (*Triticum aestivum* L.) Varieties Under Early and Late Waterlogging.” Journal of Plant Nutrition and Soil Science 185, no. 5: 622–631.

IPCC. 2023. “Summary for Policymakers.” In Climate Change 2023: Synthesis Report. Contribution of Working Groups I, II and III to the Sixth Assessment Report of the Intergovernmental Panel on Climate Change, edited by H.-J. Lee and J. Romero. Geneva, Switzerland: IPCC, 1–34.

ISO 11465. 1993. “Soil Quality – Determination of Dry Matter and Water Content on a Mass Basis – Gravimetric Method.”

Itam, M., R. Mega, S. Tadano, M. Abdelrahman, S. Matsunaga, Y. Yamasaki, K. Akashi, and H. Tsujimoto. 2020. “Metabolic and Physiological Responses to Progressive Drought Stress in Bread Wheat.” Scientific Reports 10, no. 1: 17189.

Kanehisa, M., and S. Goto. 2000. “KEGG: Kyoto Encyclopedia of Genes and Genomes.” Nucleic Acids Research 28, no. 1: 27–30.

Kang, W., and S. Kang. 2019. “On the Use of Alternative Water Use Efficiency Parameters in Dryland Ecosystems: A Review.” Journal of Ecology and Environment 43, no. 1: 24.

Keya, S. S., M. R. Islam, H. Pham, M. A. Rahman, M. Bulle, A. Patwary, M. M.-A.-R. Kanika, F. H. Hemel, T. K. Ghosh, N. Huda, Z. Hawa, M. M. Rahman, and W. Ravelombola. 2025. “Thirsty, Soaked, and Thriving: Maize Morpho-physiological and Biochemical Responses to Sequential Drought, Waterlogging, and Re-drying.” Plant Stress 15: 100722.

Komatsu, S., N. Nishiyama, and A. Diniyah. 2023. “Biochemical and Enzymatic Analyses to Understand the Accumulation of γ-aminobutyric Acid in Wheat Grown Under Flooding Stress.” Oxygen 3, no. 1: 120–132.

Lee, Y. B., W.-P. Park, M. Lee, J. Park, and K. R. Kim. 2026. “Redefining Crop Fertilization Strategies in Response to Climate Change: The Need for an Integrated Soil–nutrient–climate Approach.” Applied Biological Chemistry 69, no. 1: 13.

Lillo, C., U. S. Lea, and P. Ruoff. 2008. “Nutrient Depletion As a Key Factor for Manipulating Gene Expression and Product Formation in Different Branches of the Flavonoid Pathway.” Plant, Cell & Environment 31, no. 5: 587–601.

Lopez, G., S. H. Ahmadi, W. Amelung, M. Athmann, F. Ewert, T. Gaiser, M. I. Gocke, T. Kautz, J. Postma, S. Rachmilevitch, G. Schaaf, A. Schnepf, A. Stoschus, M. Watt, P. Yu, and S. J. Seidel. 2023. “Nutrient Deficiency Effects on Root Architecture and Root-to-shoot Ratio in Arable Crops.” Frontiers in Plant Science 13: 1067498.

Nezhadahmadi, A., Z. H. Prodhan, and G. Faruq. 2013. “Drought Tolerance in Wheat.” The Scientific World Journal 610721.

Penuelas, J., F. Coello, and J. Sardans. 2023. “A Better Use of Fertilizers is Needed for Global Food Security and Environmental Sustainability.” Agriculture & Food Security 12, no. 1: 5.

Pons, C., and C. Müller. 2022. “Impacts of Drought Stress and Mycorrhizal Inoculation on the Performance of Two Spring Wheat Cultivars.” Plants 11, no. 17: 2187.

Poorter, H., K. J. Niklas, P. B. Reich, J. Oleksyn, P. Poot, and L. Mommer. 2012. “Biomass Allocation to Leaves, Stems and Roots: Meta-analyses of Interspecific Variation and Environmental Control.” New Phytologist 193, no. 1: 30–50.

R Core Team. 2024. “R: A Language and Environment for Statistical Computing.” R Foundation for Statistical Computing, Vienna, Austria. https://www.R-project.org/.

Renziehausen, T., S. Frings, and R. Schmidt-Schippers. 2024. “’Against All Floods’: Plant Adaptation to Flooding Stress and Combined Abiotic Stresses.” The Plant Journal 117, no. 6: 1836–1855.

Ru, C., X. Hu, W. Wang, and H. Yan. 2024. “Impact of Nitrogen on Photosynthesis, Remobilization, Yield, and Efficiency in Winter Wheat Under Heat and Drought Stress.” Agricultural Water Management 302: 109013.

Ru, C., K. Wang, X. Hu, D. Chen, W. Wang, and H. Yang. 2023. “Nitrogen Modulates the Effects of Heat, Drought, and Combined Stresses on Photosynthesis, Antioxidant Capacity, Cell Osmoregulation, and Grain Yield in Winter Wheat.” Journal of Plant Growth Regulation 42, no. 3: 1681–1703.

Saha, S., and G. N. Johnson. 2025. “Divergent Effects of Successive Drought and Flooding on Photosynthesis in Wheat and Barley.” Frontiers in Plant Science 16: 1603355.

Saldanha, A. J. 2004. “Java Treeview—Extensible Visualization of Microarray Data.” Bioinformatics 20, no. 17: 3246–3248.

Schauer, N., D. Steinhauser, S. Strelkov, D. Schomburg, G. Allison, T. Moritz, K. Lundgren, U. Roessner-Tunali, M. G. Forbes, L. Willmitzer, A. R. Fernie, and J. Kopka. 2005. “GC-MS Libraries for the Rapid Identification of Metabolites in Complex Biological Samples.” FEBS Letters 579, no. 6: 1332–1337.

Schweiger, R., M. C. Baier, M. Persicke, and C. Müller. 2014. “High Specificity in Plant Leaf Metabolic Responses to Arbuscular Mycorrhiza.” Nature Communications 5: 3886.

Schweiger, R., E. Castells, L. Da Sois, J. Martínez-Vilalta, and C. Müller. 2021. “Highly Species-specific Foliar Metabolomes of Diverse Woody Species and Relationships With the Leaf Economics Spectrum.” Cells 10, no. 3: 644.

Schweiger, R., A.-M. Maidel, T. Renziehausen, R. Schmidt-Schippers, and C. Müller. 2023. “Effects of Drought, Subsequent Waterlogging and Redrying on Growth, Physiology and Metabolism of Wheat.” Physiologia Plantarum 175, no. 2: e13874.

Stallmann, J., R. Schweiger, C. A. A. Pons, and C. Müller. 2020. “Wheat Growth, Applied Water Use Efficiency and Flag Leaf Metabolome Under Continuous and Pulsed Deficit Irrigation.” Scientific Reports 10, no. 1: 10112.

Swain, R., S. Sahoo, M. Behera, and G. R. Rout. 2023. “Instigating Prevalent Abiotic Stress Resilience in Crop by Exogenous Application of Phytohormones and Nutrient.” Frontiers in Plant Science 14: 1104874.

Szabados, L., and A. Savouré. 2010. “Proline: A Multifunctional Amino Acid.” Trends in Plant Science 15, no. 2: 89–97.

Tariq, A., F. Zeng, C. Graciano, A. Ullah, S. Sadia, Z. Ahmed, G. Murtaza, K. Ismoilov, and Z. Zhang. 2023. “Regulation of Metabolites by Nutrients in Plants.” In Plant Ionomics: Sensing, Signaling and Regulation, edited by V. P. Singh and M. H. Siddiqui. Hoboken, NJ, USA: John Wiley & Sons Ltd., 1–18.

Tian, L.-X., Y.-C. Zhang, P.-L. Chen, F.-F. Zhang, J. Li, F. Yan, Y. Dong, and B.-L. Feng. 2021. “How Does the Waterlogging Regime Affect Crop Yield? A Global Meta-analysis.” Frontiers in Plant Science 12: 634898.

van den Dool, H., and P. D. Kratz. 1962. “A Generalization of the Retention Index System Including Linear Temperature Programmed Gas-liquid Partition Chromatography.” Journal of Chromatography 11: 463–471.

Voesenek, L. A. C. J., and J. Bailey-Serres. 2015. “Flood Adaptive Traits and Processes: An Overview.” New Phytologist 206, no. 1: 57–73.

Wang, J.-Y., Y.-C. Xiong, F.-M. Li, K. H. M. Siddique, and N. C. Turner. 2017. “Effects of Drought Stress on Morphophysiological Traits, Biochemical Characteristics, Yield, and Yield Components in Different Ploidy Wheat: A Meta-analysis.” In *Advances in Agronomy*: Elsevier Inc., 139–173.

Waridel, P., J.-L. Wolfender, K. Ndjoko, K. R. Hobby, H. J. Major, and K. Hostettmann. 2001. “Evaluation of Quadrupole Time-of-flight Tandem Mass Spectrometry and Ion-trap Multiple-stage Mass Spectrometry for the Differentiation of *C*-glycosidic Flavonoid Isomers.” Journal of Chromatography A 926, no. 1: 29–41.

Wouters, F. C., B. Blanchette, J. Gershenzon, and D. G. Vassão. 2016. “Plant Defense and Herbivore Counter-defense: Benzoxazinoids and Insect Herbivores.” Phytochemistry Reviews 15: 1127–1151.

Xu, H., D.-D. Jin, Y.-W. Wang, G.-H. Liu, T.-Y. Zhao, C. Liu, and L.-X. Tian. 2026. “Fertilization is More Effective in Alleviating Yield Loss of Waterlogged Crops Than Exogenous Growth Regulators: A Meta-analysis.” Frontiers in Plant Science 17: 1779872.

Xu, W., W. Zhang, Y. Yu, W. Sun, L. Yu, D. Shang, C. Xue, and Q. Zhang. 2025. “Response of Crop Photosynthetic Product Allocation Under Different Water Supply Conditions: A Global Synthetic Analysis.” Field Crops Research 333.

Yang, W., D. Qi, Y. Li, Q. Wu, J. Zhu, Y. Duan, and Q. Xiong. 2026. “Tailoring Nitrogen Input Based on Waterlogging Duration Improves Root Growth and Biomass Translocation in Cotton.” Agricultural Water Management 323: 110050.

Yurekten, O., T. Payne, N. Tejera, F. X. Amaladoss, C. Martin, M. Williams, and C. O’Donovan. 2024. “MetaboLights: Open Data Repository for Metabolomics.” Nucleic Acids Research 52, no. D1: D640–D646.

Zhang, Y., X. Chen, S. Geng, and X. Zhang. 2025. “A Review of Soil Waterlogging Impacts, Mechanisms, and Adaptive Strategies.” Frontiers in Plant Science 16: 1545912.

Zhang, Y., X.-M. Ma, X.-C. Wang, J.-H. Liu, B.-Y. Huang, X.-Y. Guo, S.-P. Xiong, and G.-X. La. 2017. “UPLC-QTOF Analysis Reveals Metabolomic Changes in the Flag Leaf of Wheat (*Triticum aestivum* L.) Under Low-nitrogen Stress.” Plant Physiology and Biochemistry 111: 30–38.

