## Supplement for "Effects of Drought, Followed by Waterlogging, on Biomass, Leaf Physiology and Chemistry of Wheat Partly Depend on Fertilisation"

**SUPPORTING INFORMATION**

**TABLE S1** Metabolites identified in wheat leaves via LC-FLD (top), GC-MS (middle) or LC-MS (bottom). Metabolite names or (for LC-MS) partial identifications (dhex, deoxyhexosyl; hex, hexosyl; pen, pentosyl) along with their abbreviations in brackets are shown. Metabolites, for which identification was based on a reference standard, are presented in bold. Names of acids are given in deprotonated form. In addition, retention parameters (LC-FLD, LC-MS: retention times; GC-MS: arithmetic retention indices) and mass-to-charge ratios (*m*/*z*; GC-MS: characteristic *m*/*z*; LC-MS: accurate *m*/*z* of main ion with ion type in MS mode in first line, fragments in MS/MS mode with dominant ones in bold in second line) are provided. Due to co-elution, orientin and isoorientin could not be distinguished.

| Analytical platform and metabolite | Retention parameter | *m/z* |
|---|---|---|
| **LC-FLD** | | |
| **Amino acids** | | |
| **aspartate** [ASP] | 1.4 min | |
| **glutamate** [GLU] | 1.9 min | |
| **asparagine** [ASN] | 3.1 min | |
| **serine** [SER] | 3.3 min | |
| **glutamine** [GLN] | 3.7 min | |
| **histidine** [HIS] | 3.9 min | |
| **homoserine** [HOS] | 4.1 min | |
| **glycine** [GLY] | 4.2 min | |
| **threonine** [THR] | 4.4 min | |
| **citrulline** [CIT] | 4.6 min | |
| **β-alanine** [βALA] | 5.0 min | |
| **arginine** [ARG] | 5.2 min | |
| **alanine** [ALA] | 5.6 min | |
| **γ-aminobutyrate** [GABA] | 5.9 min | |
| **tyrosine** [TYR] | 7.2 min | |
| **valine** [VAL] | 9.7 min | |
| **methionine** [MET] | 10.0 min | |
| **tryptophan** [TRP] | 11.3 min | |
| **phenylalanine** [PHE] | 11.9 min | |
| **isoleucine** [ILE] | 12.2 min | |
| **leucine** [LEU] | 13.1 min | |
| **proline** [PRO] | 16.8 min | |
| **GC-MS** | | |
| **Carbohydrates** | | |
| **fructose** [Frc] | 1859\|1869 | 217\|277\|364\|335\|307 |
| **glucose** [Glc] | 1883\|1901 | 319\|229\|343\|305\|160 |
| **sucrose** [Suc] | 2613 | 451\|361\|319\|157\|437 |
| **Organic acids** | | |
| **succinate** [SucA] | 1311 | 172\|147\|262\|129\|247 |
| **glycerate** [GlyA] | 1324 | 189\|307\|205\|133\|292 |
| **itaconate** [ItaA] | 1338 | 215\|133\|147\|230\|259 |
| **malate** [MalA] | 1483 | 245\|335\|307\|217\|233 |
| **aconitate** [AcoA] (*cis*- and/or *trans*-) | 1745 | 285\|375\|211\|215\|229 |
| **citrate** [CitA] | 1810 | 375\|211\|183\|257\|273 |
| **Cyclic polyol** | | |
| ***myo*-inositol** [Ino] | 2074 | 265\|318\|191\|507\|305 |
| **Inorganic acid** | | |
| **phosphate** [PhoA] | 1264 | 299\|211\|283\|225\|314 |
| **LC-MS** | | |
| **Benzoxazinoids** | | |
| **DIMBOA glucoside** [DIMBOA glc] | 7.62 min | 372.09349 $[M-H]^{-}$<br>192\|164\|**149** |
| **HDMBOA glucoside** [HDMBOA glc] | 10.37 min | 432.11484 $[M+HCOOH-H]^{-}$<br>**194**\|164\|149 |
| **Flavonoids** | | |
| luteolin *C*-hex, *C*-pen [Lut *C*-hex *C*-pen] | 9.23 min | 579.13554 $[M-H]^{-}$<br>519\|**489**\|**459**\|429\|399\|369 |
| **(iso)orientin** [(Iso)Ori] | 10.01 min | 447.09334 $[M-H]^{-}$<br>429\|411\|**357**\|339\|**327**\|297\|285 |
| **schaftoside** [Schaf] | 10.16 min | 563.14090 $[M-H]^{-}$<br>545\|503\|**473**\|**443**\|383\|353 |

<table>
<tr><td>luteolin C-hex, 2”-O-dhex [Lut C-hex O-dhex]</td><td>10.25 min</td><td>593.15136 [M-H]$^-$<br>575|503|473|429|357|339|327|309</td></tr>
<tr><td>isovitexin [Isovit]</td><td>11.46 min</td><td>431.09825 [M-H]$^-$<br>353|341|323|311|283</td></tr>
<tr><td>apigenin C-hex, 2”-O-dhex [Api C-hex O-dhex]</td><td>11.57 min</td><td>577.15633 [M-H]$^-$<br>457|413|341|323|311|293</td></tr>
<tr><td>apigenin 6,8-di-C-pen [Api di-C-pen]</td><td>11.82 min</td><td>533.13006 [M-H]$^-$<br>515|473|443|425|383|353</td></tr>
<tr><td>chrysoeriol C-hex [Chr C-hex]</td><td>12.29 min</td><td>461.10906 [M-H]$^-$<br>443|371|353|341|298</td></tr>
<tr><td>chrysoeriol C-hex, 2”-O-dhex [Chr C-hex O-dhex]</td><td>12.33 min</td><td>607.16718 [M-H]$^-$<br>487|443|371|353|341|323</td></tr>
<tr><td>tricin di-O-hex [Tri di-O-hex]</td><td>12.70 min</td><td>653.17228 [M-H]$^-$<br>329</td></tr>
<tr><td>tricin O-hex, O-dhex [Tri O-hex O-dhex]</td><td>14.07 min</td><td>637.17749 [M-H]$^-$<br>329</td></tr>
<tr><td>tricin [Tri]</td><td>20.29 min</td><td>329.06670 [M-H]$^-$<br>314|299|271</td></tr>
</table>

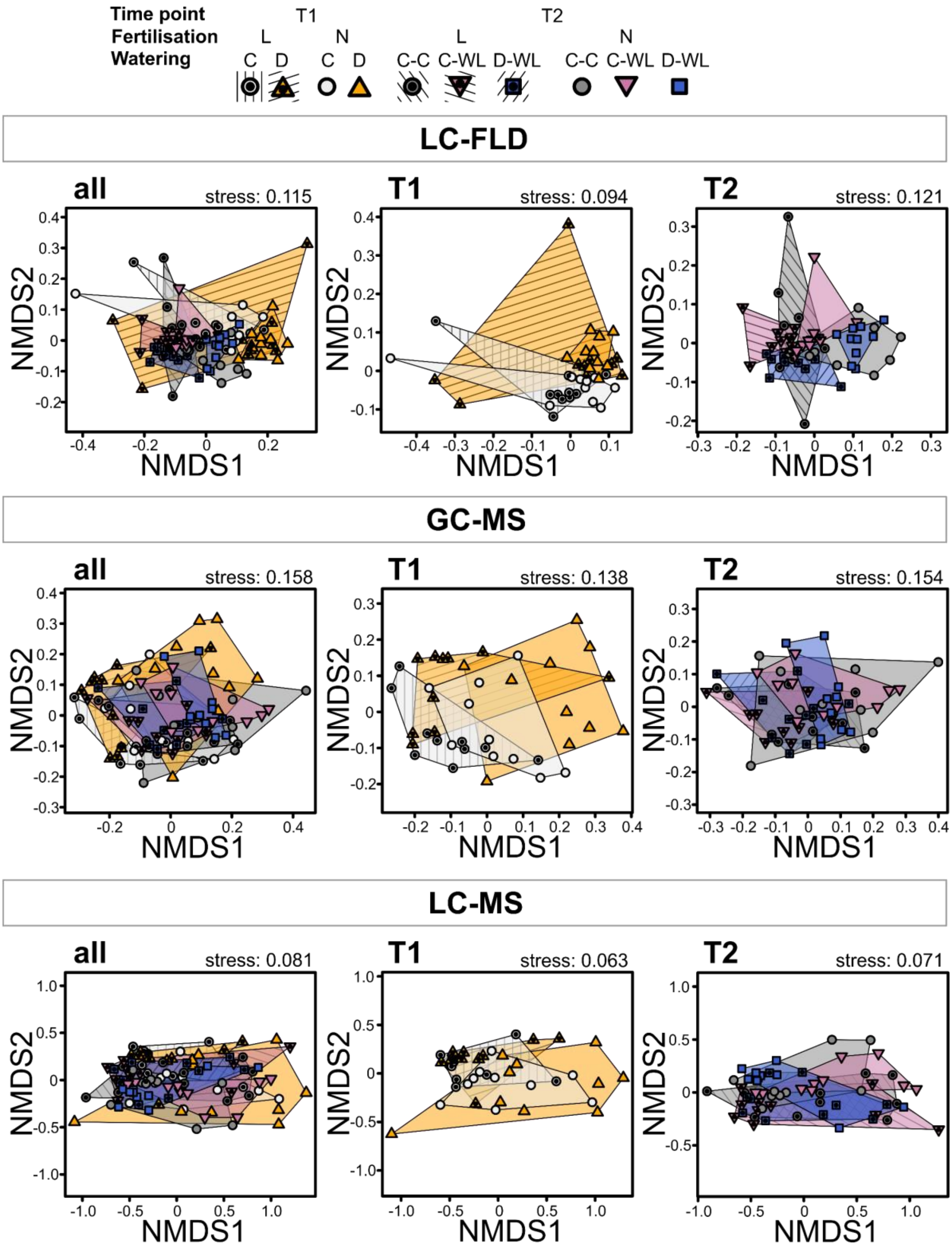


**FIGURE S1** Metabolite patterns of *Triticum aestivum* leaves, shown as non-metric multidimensional scaling (NMDS) plots. Plants were subjected to different watering regimes (well-watered control, C; drought, D) in the drought phase, while in the subsequent waterlogging phase they were well-watered (C) or waterlogged (WL). Two fertilisation treatments were applied (low, N; normal, N). Leaves were harvested at two time points (end of drought phase, T1; end of waterlogging phase, T2). Separate analyses were done for amino acids (LC-FLD, first row), further primary metabolites (GC-MS, second row) and for metabolic fingerprints mainly containing specialised metabolites (LC-MS, third row). NMDS plots including all samples are shown on the left, while time point-specific analyses are presented in the middle (T1) and right (T2). Stress values of the NMDS analyses are provided at the top right. Scores are given as individual data points (containing a small black circle for the L treatment group). Groups are surrounded by convex hulls (hatched for the L treatment); *n*=9-10.

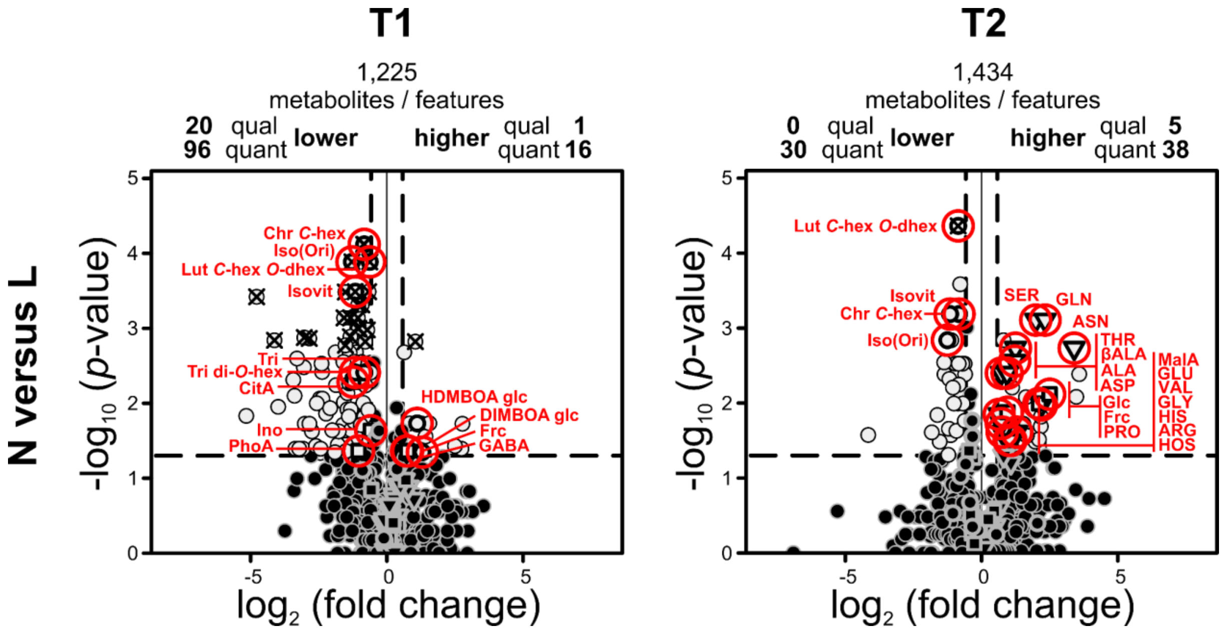


**FIGURE S2** Modulation of metabolites and metabolic features in *Triticum aestivum* leaves by fertilisation, shown as volcano plots. Plants were well-watered and two fertilisation treatments (low: L; normal: N) were applied; leaves were harvested at two time points (T1: left; T2: right). For each group comparison (N versus L within time points), *p*-values (Mann-Whitney *U*-tests, $-\log_{10}$) and mean fold changes ($\log_2$) of metabolites/ features occurring in both groups are plotted. Grey symbols represent modulated metabolites/ features outside the dashed cutoff lines (horizontal: corresponding to *p*-value of 0.05; vertical: corresponding to fold changes of 0.67 and 1.5), with those on the left being lower and those on the right being higher in the normal fertilisation than in the low fertilisation group. Non-modulated metabolites/ features are shown in black. Symbol types indicate analytical platforms (triangles: LC-FLD, amino acids; squares: GC-MS, further primary metabolites; circles: LC-MS, mainly specialised metabolites). (Partially) identified metabolites are surrounded by thicker lines and labeled in red if they were quantitatively modulated (full names: Table S1); there were no identified metabolites among the qualitatively modulated features. Metabolites/ features that were significant after Benjamini-Hochberg correction of *p*-values are indicated with crosses. The numbers of qualitatively and quantitatively modulated features are given at the top. Based on *n*=9-10.

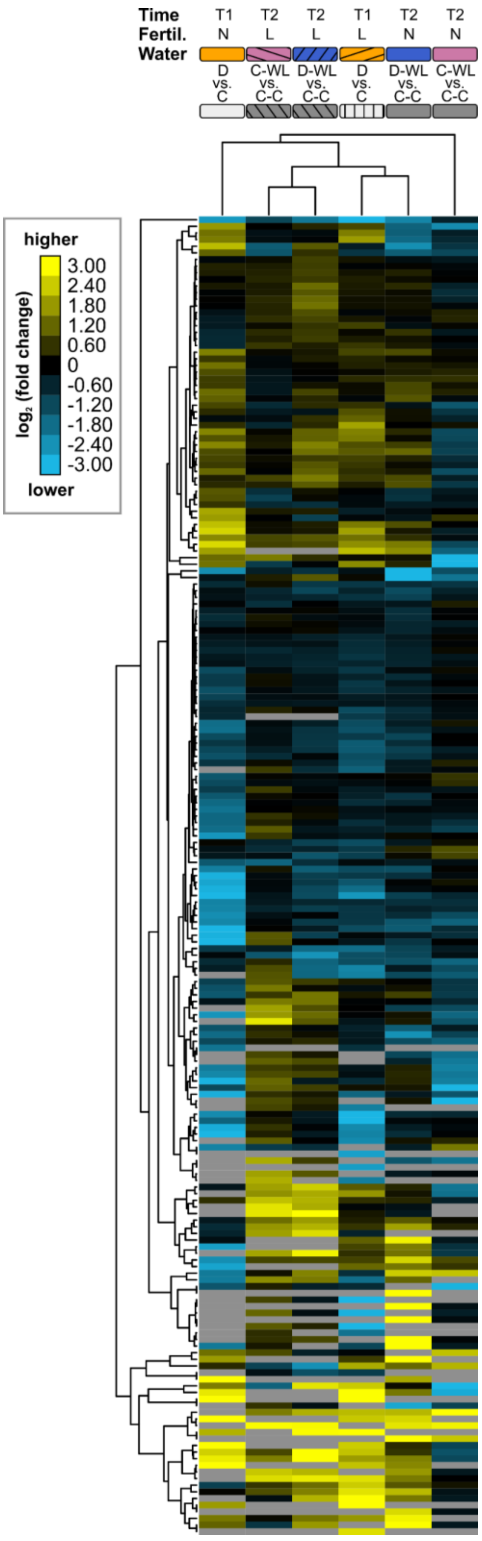


**FIGURE S3** Modulation of metabolites and metabolic features in *Triticum aestivum* leaves by drought and/ or waterlogging, shown as cluster heatmap. Plants were subjected to different watering regimes (well-watered control, C; drought, D) in the drought phase, while in the subsequent waterlogging phase they were well-watered (C) or waterlogged (WL). Two fertilisation treatments were applied (low, N; normal, N). Leaves were harvested at two time points (end of drought phase, T1; end of waterlogging phase, T2). Fold changes of those 199 metabolites/ features were included, which were quantitatively modulated ($p$-value < 0.05 in Mann-Whitney $U$-test; $\log_2$-scaled fold changes < 0.58 or > 0.58) in at least one group comparison. Yellow and blue indicate higher and lower concentrations in the drought/ waterlogging compared to the control group, respectively, while features that did only occur in one group are indicated in grey; based on $n$=9-10.
